# A Sample to Results Workflow for Compositional Analysis of Multiplexed Amplicon Sequencing Experiments

**DOI:** 10.64898/2026.07.28.741237

**Authors:** Alexa Bennett, Ryan Moore, Craig W. Herbold, Thomas E. Hanson

## Abstract

Microbial communities play key roles in the transformation and cycling of elements ranging from required macronutrients to toxic metalloids. Next-generation sequencing has been applied across multiple ecosystems to probe the interplay of microbial community structure and functional potential with respect to elemental cycling. Shotgun metagenomics collects marker gene sequences without amplification and is costly for large numbers of samples and deep coverage. Conversely, amplicon sequencing of taxonomic marker genes, e.g. 16S and 18S rRNA, is cost-effective for large numbers of samples, but provides limited functional insight. A middle ground between the two approaches is needed to analyze community structure and functional potential within a sample while remaining cost-effective with high throughput. To address this need, we developed a standardized workflow for multiplexed amplicon sequencing from sample collection through data analysis for diverse sample types, including freshwater, sediments, and soils, that produces data and publication-ready figures for multiple taxonomic and functional genes for carbon, nitrogen, phosphorus, sulfur, and arsenic cycling for each sample analyzed. The workflow’s utility was shown by analyzing 11 taxonomic and functional gene amplicons sequenced from 25 samples with high technical replicate similarity. The workflow is named *<u>CAMASE</u>* for *<u>C</u>*ompositional *<u>A</u>*nalysis of *<u>M</u>*ultiplex *<u>A</u>*mplicon *<u>S</u>*equencing *<u>E</u>*xperiments. This proof-of-concept shows that CAMASE economically produces standard amplicon sequencing outputs (ASV/OTU counts and taxonomy, PCA, and relative abundance plots) for hundreds of amplicon by sample combinations and provides specific recommendations for implementation.

**GRAPHICAL ABSTRACT:** 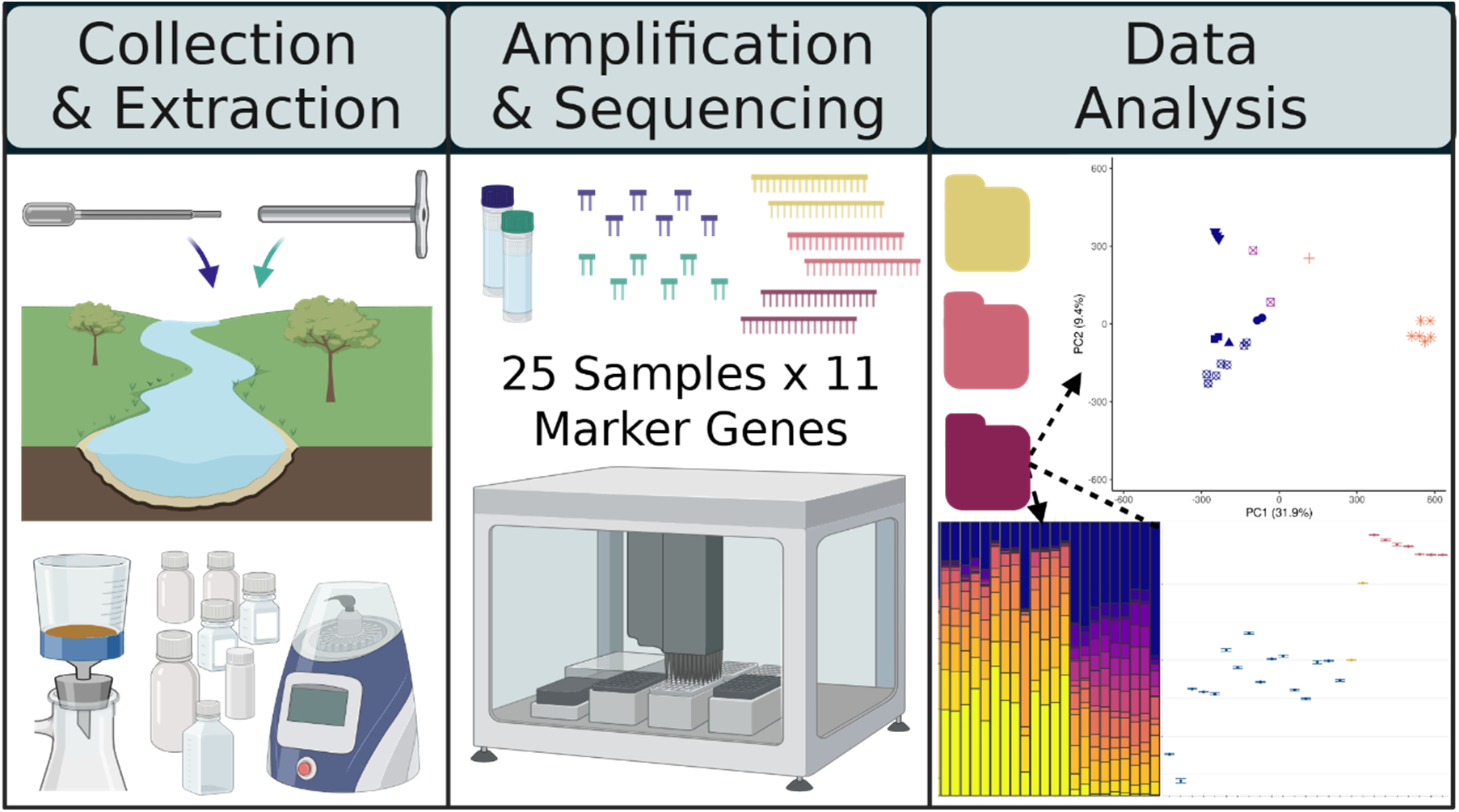

Samples are collected in a preservative and material collected on filters prior to DNA extraction. Target gene amplicons are produced in parallel with internal barcodes enabling sequencing in a single run followed by compositional data analysis. All wet lab protocols, code markdowns, and templates for required metadata files are available at https://hansonlabgit.dbi.udel.edu/aprange/CAMASE. Created in BioRender. Bennett, A. (2026) https://BioRender.com/ymnojt0

## INTRODUCTION

Microbial communities drive biogeochemical cycling of carbon, nitrogen, phosphorus, and sulfur. Their composition is crucial for understanding the distribution of elemental species and fully describing an environment or ecosystem [1, 2]. For example, the nitrogen cycle contains transformations between gaseous (nitrogen gas, nitrous oxide), organic (urea, amino acids), and mineral species (ammonium, nitrate, nitrite), all of which can be oxidized or reduced by specific pathways encoded by the genomes of multiple groups of bacteria, archaea, and fungi [3]. Nitrogen, as ammonium and nitrate, is commonly the limiting nutrient in terrestrial primary productivity underscoring the importance of microbial biogeochemistry [4, 5]. Microbes also cycle and speciate toxic elements, e.g., arsenic [6]. Environmental microbiome surveys are used to connect community composition with functional potential, but the appropriate strategy depends on whether the goal is broad taxonomic profiling, targeted functional gene detection, or genome-resolved inference [7].

Low-cost high-throughput protocols targeting phylogenetic markers, e.g. 16S and 18S rRNA genes have revolutionized our ability to determine “who” is present in a microbiome [8, 9]. Computational approaches exist that attempt to infer functional potential from 16S rRNA amplicon sequence data, but they have limited predictive power on environmental samples [10, 11]. Alternatively, metagenomic sequencing captures functional gene information in a microbiome. However, there are inherent trade-offs between sequencing effort, cost, and gene/genome coverage depth. Metagenomics can achieve deep coverage of moderately abundant community members, but only with significant effort and cost. Incomplete coverage of functional genes in low-abundance, but functionally important, community members can occur, particularly in complex microbial communities that have many low abundance members that are collectively called the rare biosphere [7].

Here we describe a middle ground between single amplicon surveys and metagenomics named *<u>CAMASE</u>* for *<u>C</u>*ompositional *<u>A</u>*nalysis of *<u>M</u>*ultiplex *<u>A</u>*mplicon *<u>S</u>*equencing *<u>E</u>*xperiments. CAMASE provides detailed and sensitive analysis of both functional and phylogenetic gene amplicons. CAMASE is based on a flexible two-step barcoding approach [12]. It extends this approach by standardizing sample collection, preservation, DNA isolation methods, and incorporating liquid handling to minimize batch effects on sequence library generation. The computational pipeline primarily uses Python and R for ease of use by non-bioinformaticians. CAMASE implements data analyses that recognize the inherently compositional nature of microbiome amplicon sequencing data [13, 14]. Detailed standard operating procedures (SOPs), robotics programs, calculation templates, and data analysis scripts are publicly accessible at: https://hansonlabgit.dbi.udel.edu/aprange/CAMASE.

## MATERIALS AND METHODS

### Environmental Sample Collection & DNA Extraction

CAMASE (Fig. 1) begins with sample collection. Environmental samples (freshwater, sediment, and soil) were collected with sterile disposable pipettes or autoclaved reusable metal corers and immediately mixed 1:1 volume:volume with 20 mL DMSO-EDTA-Saturated Salt (DESS) [15] for transport to the laboratory, where they were stored at 4°C until purification. Twenty environmental samples were collected for this proof-of-concept study with three soil and two freshwater samples analyzed twice starting with PCR amplification as technical replicates for a final count of twenty-five samples (Table 1, Supplemental File 1).

**Figure 1.**
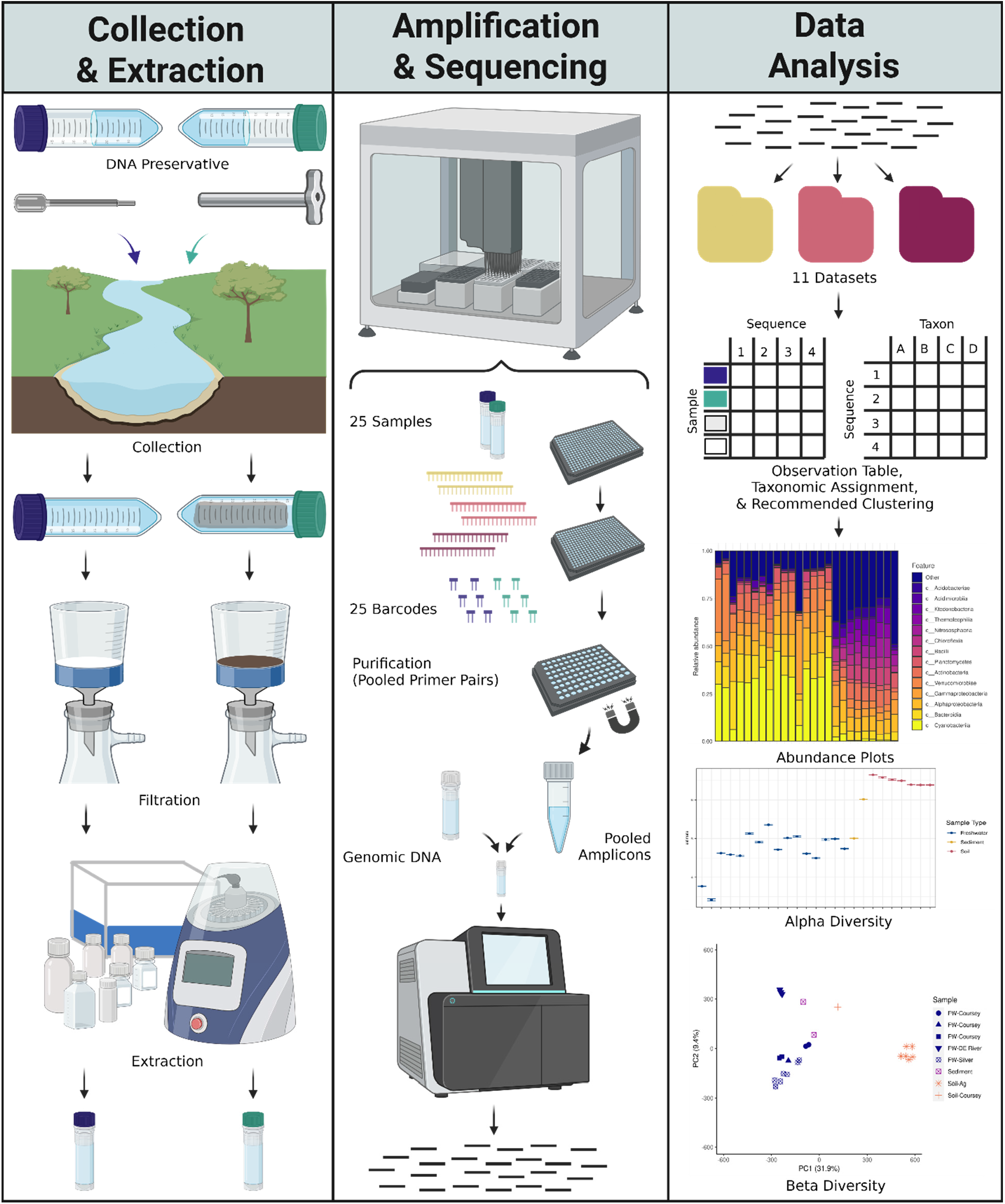
Overview of CAMASE workflow. For simplicity, the workflow overview follows two samples, a water sample in blue and a soil sample in green, with three gene-specific primer sets: yellow, pink, and red. The workflow includes sample collection with a transfer pipette or metal corer into a DNA preservative. Samples are filtered, followed by DNA extraction. A robotic liquid handler is utilized for a 2-step PCR approach, which amplifies a gene-specific target and then adds a sample-identification barcode. Samples are pooled within gene-specific targets and purified with a magnetic bead protocol. Purified gene-specific amplicon pools are combined with the addition of non-amplified genomic DNA. The final pool of DNA is then processed for a single MiSeq library preparation and analysis. Reads from the MiSeq analysis are separated by dataset (gene-specific target). Within each dataset, samples are demultiplexed and processed to create a sequence or operational taxonomic unit (OTU) table and taxonomic reference table. Within each dataset, relative abundance, alpha diversity estimates, and beta diversity are visualized. Created in BioRender. Bennett, A. (2026) https://BioRender.com/4ppdwpx

**Table 1.** The CAMASE version 1 amplicon panel genes and gene product functions.

| Gene(s)/Dataset<sup>1</sup> | Process | Gene Product |
| --- | --- | --- |
| 16S rRNA | Bacterial and Archaeal community structure | 16S ribosomal RNA |
| 18S rRNA | Eukaryotic community structure | 18S ribosomal RNA |
| *arrA* | Arsenic resistance and transformation | Respiratory arsenate reductase catalytic subunit |
| *dsrB* | Sulfur reduction and oxidation | Dissimilatory type sulfite reductase subunit beta |
| *gcd* | Inorganic phosphorus dissolution | Quinoprotein glucose dehydrogenase |
| *mcrA* | Methanogenesis-methane production | Methyl coenzyme-M reductase subunit |
| *narG* | Nitrate reduction to nitrite | Nitrate reductase subunit |
| *nirK* | Denitrification - nitrite reduction to N<sub>2</sub> | Nitrite reductase |
| *nrfA* | Dissimilatory nitrate reduction to ammonium | Nitrite reductase |
| *phoD_X* | Mineralization of organic phosphorus | Alkaline phosphatase |
| *phoN* | Mineralization of organic phosphorus | Acid phosphatase |
| *pmoA* | Methanotrophy-methane consumption | Particulate methane monooxygenase subunit |

1 - No amplification was observed for *gcd* or *phoN* genes using the primers selected (Table S2)

DNA extraction was initiated by hand shaking samples followed by vacuum filtration onto a 0.10 µm PVDF hydrophilic membrane (Durapore®). A maximum of 500 mg of biomass and filter was used for extraction by the FastDNA™ Spin Kit for soil (MP Biomedical) following manufacturer instructions with homogenization in a FastPrep-24™ Classic (MP Biomedical) for 40 sec at 6.0 m/s.

DNA was quantified with Qubit™ dsDNA HS Assay Kit (Life Technologies) and a Qubit® 2.0 Fluorometer system (Life Technologies) or Quant-iT™ dsDNA High-Sensitivity Assay Kit (Life Technologies) and a SpectraMax i3x plate reader (Molecular Devices) with 485 nm excitation and 530 emission wavelengths. Extracts greater than 0.5 ng DNA/µL were diluted to 0.5 ng/µL.

### PCR Amplification

Thirteen gene-specific primer sets were synthesized (Eurofins). This panel includes primers for 16S and 18S rRNA, and functional genes related to carbon, nitrogen, phosphorous, sulfur, and arsenic (Table 1). While most primer sets consist of single degenerate forward and reverse primer sequences (Table S1), the *dsrB* gene was targeted using a mix of eight forward and eight reverse degenerate primer sequences (Table S2). Two primer pairs, *phoD* and *phoX*, were combined and treated as a single primer set, *phoD_X* (Table S2).

Three control types were constructed. The positive control was constructed from an unbalanced mock community of purified genomic DNAs obtained from the American Type Culture Collection (Table S3). The negative control was molecular grade water (Fisher Scientific). The process control consisted of DESS preservative filtered and extracted as described for samples. The positive and process controls are sequenced for each run while the negative control was used in fragment analysis, but was not sequenced.

Template DNA was amplified in two steps [12]. The first PCR reaction contained gene-specific primers (Table S1 & Table S2) with the head sequence [5′-GCTATGCGCGAGCTGC-3′]. The second PCR reaction primers contain the head sequence at the 3’-end preceded by a unique 8bp barcode sequence, enabling sample identification during data analysis. Barcodes were selected in sets of 28 with a minimum Hamming distance of three between any two barcodes per sequencing run [16, 17]. Alterations to the original protocol [12] were: ⅰ) single gene-specific PCR reactions, ⅱ) 20 μL reaction volumes for both steps, ⅲ) 2 µL of template, ⅳ) Platinum™ SuperFi II PCR Master Mix (Invitrogen™), and ⅴ) 60 ℃ annealing. PCR conditions are provided in Supplemental Material (Table S4). PCR Master Mix, water, and primers were premixed, aliquoted, and stored at −20 ℃ [18]. Amplicon concentrations were measured by Quant-IT, as described above. Amplicon size and quality were assessed by fragment analysis (Advanced Analytical Technologies 5200 Fragment Analyzer, 33cm array).

### Amplicon Cleanup, Pooling, and Sequencing

Amplicons were diluted to 5 ng/µL where necessary. Equal volumes of each reaction for a given amplicon were mixed, then purified with SparQ PureMag Beads (QuantaBio). Purified reaction pools were combined at 0.39 nM for *arrA*, *dsrB*, *mcrA*, *nirK*, *nrfA*, and *pmoA*, and 0.77 nM for 16S rRNA, 18S rRNA, *narG*, and *phoD_X.* Sheared (Diagenode One, Diagenode Epigenetics) genomic DNA fragments of 300-500 bp (BluePippin, Sage Science) were added at 0.60 nM to constitute ∼10% of the final pool to add sequence complexity required for Illumina sequencing. PCR reaction assembly, quantification assays, amplicon purification, and pooling were performed on a SciClone G3 NGSx liquid handling workstation (Revvity).

The final pool was transferred to the University of Delaware DNA Sequencing & Genotyping Center. MiSeq library preparation was performed with the SparQ DNA Library Prep Kit (QantaBio) according to the protocol without the fragmentation step and sequenced using Illumina with MiSeq Reagent Kit v3. The same library was analyzed in two separate MiSeq runs, one 2 x 300 cycle run and one 2 x 330 cycle run.

### Data Analysis

A brief description of the data analysis pipeline follows. A detailed tutorial with code markdowns is available at: https://hansonlabgit.dbi.udel.edu/aprange/CAMASE. Raw Illumina reads were parsed into datasets, which contain all sequences produced by a gene-specific primer set. Samples within datasets were demultiplexed by sample-identification barcode, followed by removal of 5’-end primer sequences (barcode, head sequence, gene-specific primer) using the same Python script as Herbold et al. [12] with Python 3.10.10 and QIIME2 2021.2 [19]. Amplicon sequence variants (ASVs) were produced for each dataset with independent error learning and denoising with DADA2 v1.21.0 [20] per “DADA2 pipeline Tutorial (1.16),” trimmed where the median quality score dropped below Q30 with FASTX-toolkit v0.0.14 [21] and summarized using in-house BASH and Python scripts. Following trimming, reads 1 and 2 for short amplicons (< 450 bp) were merged, while large amplicons (> 450 bp) were concatenated after trimming.

ASVs were classified with the RDP classifier (Q. Wang et al. 2007) using curated taxonomic databases. Taxonomic databases were constructed for the SSU rRNA datasets from SILVA [22–24], while functional gene databases used FunGene [25] and NCBI [26, 27]. An in-house modification of RESCRPt [28] was used to retrieve taxonomic identifications, remove redundancies and low-quality sequences, and trim to the amplicon target region. The RESCRPt modifications ensured only assigned taxonomies were returned and handled multiple gene copies within single assembly accession numbers. The full curation workflow and curated FASTA classifiers are at: https://hansonlabgit.dbi.udel.edu/aprange/simplified_taxonomic_curation

ASVs were aligned and clustered with DECIPHER v2.22 [29] from 97% to 75% similarity using the “complete” method. When memory became a limitation for large dataset alignment, a chained guide tree was used as described [30]. Clusters were collapsed into operational taxonomic units (OTUs) and retained for downstream processing if observed in two or more samples.

Beta diversity was explored by plotting relative abundance values for ASVs, ASVs collapsed by taxon rank, and OTU using the R package FeatureTable (v0.0.11) [31]. Abundance of the OTU clusters produced by DECIPHER and ASVs collapsed by taxon rank were used for alpha diversity analysis with DivNet [14]. Calculations were performed using sample names as the only covariate, tuning = “careful”, and network = “diagonal.” For datasets without an OTU common to all samples, an OTU with the highest median number of observations amongst the most common OTUs within a dataset was used at the base OTU. The remaining parameters were run with the default settings. Shannon diversity estimates from DivNet were then visualized with ggplot2 [32].

Principal component analysis (PCA) was performed on the Aitchison distances using the R package Vegan (v2.6-6.1). Features were first transformed with zCompositions square root Bayesian-Multiplicative replacement, z.warning = 1, z.delete = FALSE, followed by centered-log ratio (clr) transformation [33]. The Euclidean distance of the transformed data is equivalent to the Aitchison distance [34].

### Assessing CAMASE Reproducibility

The reproducibility of the workflow was assessed by analyzing the technical replicates. After zero replacement, a distance matrix was constructed for only the technical replicates using vegdist() from the Vegan package. The distance matrix was assessed by Mantel test with mantelTest() and mantelPower() in the Biotools package with an effect size of 1. PCA was performed and visualized within the context of all samples to check that replicates clustered together.

## RESULTS

CAMASE was evaluated with a panel of 13 taxonomic and functional gene amplicons (Table 1) and 25 samples collected from freshwater, sediment, and soil environments (Table 2). Amplicons were selected with input from local colleagues and provide data on 1) community taxonomy, 2) microbial cycling of carbon, nitrogen, phosphorous, and sulfur, and 3) reduction of the toxic metalloid arsenic. The data presented summarize CAMASE performance in terms of amplicon success, sequencing quality, sequence retention, technical reproducibility and illustrate the standard outputs of compositional microbiome analysis.

**Table 2.** Samples analyzed in this study.

| Name<sup>1</sup> | DNA<sup>2</sup> | Final Read Pairs per Dataset<sup>3</sup> |  |  |  |  |  |  |  |  |  |  |
| --- | --- | --- | --- | --- | --- | --- | --- | --- | --- | --- | --- | --- |
|  |  | 16S | 18S | *arrA* | *dsrB* | *mcrA* | *narG* | *nirK* | *nrfA* | *phoD* | *phoX* | *pmoA* |
| Positive_Control | NA | 116055 | 58589 | 420 | 6119 | 32988 | 8787 | 28031 | 47405 | 79 | 5678 | 8734 |
| Process_Control | NA | 1813 | 1627 | 10 | 4 | 1 | 3 | ND | 10 | 1 | 1 | 9 |
| FW-Coursey Pond LT-beg-1 | Optimal | 159025 | 188482 | 6122 | 17975 | 15749 | 6830 | 5284 | 25060 | 15789 | 10528 | 36002 |
| FW-Coursey Pond LT-beg-2 | Sub-optimal | 127376 | 184594 | 4424 | 14670 | 9111 | 5464 | 2564 | 17642 | 5729 | 6648 | 24725 |
| FW-Coursey Pond LT-end-1 | Optimal | 150026 | 173433 | 4177 | 3993 | 5165 | 3368 | 6878 | 6816 | 8155 | 8223 | 24864 |
| FW-Coursey Pond MOD-1 | Sub-optimal | 117487 | 104511 | 152 | 113 | 288 | 43 | 93 | 235 | 385 | 1 | 174 |
| FW-Coursey Pond MOD-2 | Optimal | 27015 | 168502 | 99 | 98 | 6 | 73 | 6 | 20 | 78 | 2 | 26 |
| FW-DE River-1 | Sub-optimal | 153041 | 209176 | 2044 | 2309 | 2 | 671 | 629 | 7021 | 1323 | 740 | 6289 |
| FW-DE River-2 | Sub-optimal | 117083 | 295139 | 2151 | 1940 | 222 | 1810 | 1566 | 10246 | 2115 | 1060 | 8693 |
| FW-DE River-3 | Sub-optimal | 45854 | 112781 | 4978 | 6708 | 3783 | 3276 | 4353 | 14085 | 1503 | 326 | 12145 |
| FW-Silver Lake-1 | Optimal | 175398 | 185574 | 1832 | 2605 | 2342 | 791 | 693 | 3638 | 3745 | 4075 | 34551 |
| FW-Silver Lake-2* | Sub-optimal | 116915 | 227045 | 1500 | 2261 | 975 | 899 | 3856 | 6495 | 2444 | 2463 | 16333 |
| FW-Silver Lake-3* | Sub-optimal | 118602 | 175086 | 1290 | 1125 | 1413 | 890 | 2972 | 5442 | 1459 | 1592 | 11325 |
| FW-Silver Lake-4 | Optimal | 62691 | 107016 | 6784 | 7122 | 2537 | 2845 | 4758 | 25233 | 1145 | 1227 | 9645 |
| FW-Silver Lake-5 | Sub-optimal | 110519 | 163059 | 59 | 216 | 3 | 106 | 194 | 149 | 633 | 448 | 7469 |
| FW-Silver Lake-6* | Sub-optimal | 50809 | 139060 | 309 | 562 | 642 | 302 | 687 | 657 | 1034 | 531 | 9794 |
| FW-Silver Lake-7* | Sub-optimal | 94557 | 118058 | 524 | 861 | 213 | 730 | 584 | 688 | 897 | 345 | 4716 |
| FW-Silver Lake-8 | Optimal | 102716 | 155928 | 392 | 2521 | 2461 | 807 | 2348 | 3661 | 4770 | 6071 | 19753 |
| Sediment-DE River-1 | Sub-optimal | 15543 | 4478 | 16 | 30 | 2 | 1 | 2 | 555 | 349 | 213 | 10 |
| Sediment-DE River-2 | Sub-optimal | 90711 | 56337 | 1453 | 275 | 371 | 696 | 204 | 8984 | 1389 | 2454 | 692 |
| Soil-Agricultural-1* | Optimal | 111129 | 106309 | 3995 | 1122 | 99 | 6898 | 7387 | 18710 | 7577 | 3117 | 1149 |
| Soil-Agricultural-2* | Optimal | 67084 | 84550 | 4544 | 1401 | 3 | 5225 | 5399 | 21773 | 3109 | 1949 | 987 |
| Soil-Agricultural-3* | Optimal | 128660 | 115253 | 2066 | 1073 | 7 | 2519 | 4305 | 7332 | 2731 | 1129 | 868 |
| Soil-Agricultural-4* | Optimal | 66721 | 68442 | 1766 | 860 | 7 | 2746 | 2782 | 2595 | 1136 | 592 | 748 |
| Soil-Agricultural-5* | Optimal | 129984 | 138067 | 4702 | 2374 | 8 | 2370 | 7602 | 32969 | 24814 | 5810 | 1984 |
| Soil-Agricultural-6* | Optimal | 70028 | 89325 | 2931 | 2334 | 13 | 9015 | 9411 | 12316 | 5642 | 2622 | 1205 |
| Soil-Coursey Landing-1 | Optimal | 73302 | 169261 | 53070 | 43353 | 707 | 24767 | 38429 | 32560 | 44930 | 6589 | 11205 |

1 - Name used in figures, asterisks indicate technical replicate pairs, e.g. samples analyzed in parallel from the gene-specific PCR
2 - DNA mass in gene-specific PCR: Optimal = 1.0 ng, Sub-optimal < 1.0 ng, NA = Not applicable
3 – Sum of final read pairs produced for a given dataset and sample. ND = Not detected

**Table 3.** Mantel test results of 95% Similarity OTUs. The input was the Aitchison distance calculated between technical replicate sample pairs (asterisks in. Table 1).

| **Dataset** | **r** | **p-value** | **power** |
| --- | --- | --- | --- |
| 16S | 0.991 | 0.001 | 0.405 |
| 18S | 0.997 | 0.001 | 0.395 |
| *arrA* | 0.539 | 0.190 | 0.775 |
| *dsrB* | 0.873 | 0.001 | 0.518 |
| *mcrA* | 0.926 | 0.001 | 0.503 |
| *narG* | 0.936 | 0.001 | 0.371 |
| *nirK* | 0.938 | 0.001 | 0.498 |
| *nrfA* | 0.980 | 0.001 | 0.506 |
| *phoD* | 0.991 | 0.001 | 0.414 |
| *phoX* | 0.984 | 0.001 | 0.425 |
| *pmoA* | 0.922 | 0.001 | 0.502 |

### Amplicon Panel Assessment

Gene-specific PCR was performed with a range of DNA concentrations, 0.025-3.59 ng/µL for the positive control and 0.025-5.02 ng/µL for a soil DNA extract. The optimal DNA template mass (1.0 ng) was selected by a marked increase in amplicon concentration compared to the negative control amongst the functional genes with no inhibition of 16S and 18S rRNA amplification. CAMASE targeted 13 genes (Table 1), but no *gcd* and *phoN* amplicons were observed (Table S5). Testing at a reduced annealing temperature (50.5℃) also produced no *gcd* and *phoN* amplicons (data not shown) and they were omitted from further analysis. No *phoD* amplicon in the correct size range was observed in the positive control, despite the mixture containing 800 copies of *phoD* from *Bacillus subtilis*. Soil amplified with *phoD_X* contained multiple amplicons, one within the range expected for *phoX*, but none within the expected range for *phoD* (Table S5). Even so, the presence of *phoD* sequences in the amplicons was validated as described in the next section.

### Sequencing Quality and Retention

Sequence retention is the amount of data that passes CAMASE sequence parsing and quality filtering and was evaluated by tallying final read pair counts, the number of paired reads unambiguously assigned to a dataset from a given sample. CAMASE amplicons range in size from ∼300-700 nt with adapters and barcodes (Tables S1 & S2). MiSeq paired end reads were trimmed based on a quality score of 30 (Q30). Reads for shorter amplicons, e.g. 16S rRNA and *dsrB*, can be merged while longer amplicon sequences, e.g. *nirK*, need to be concatenated after quality trimming. Either end of the amplicon can be sequenced in read 1 or read 2, resulting in four files per amplicon that need trimming. Each of the four files can be trimmed individually (Trim per File), or the shortest Q30 length of a dataset can be set as the trim length for all files (Trim per Dataset Minimum). Trim per Dataset Minimum resulted in an average loss of 110 bp in concatenated amplicon length (Fig. 2A, open vs. closed circles). The negative trend of read pair length in longer amplicons was due to shorter trimmed read 2 lengths (Fig. 2A, coefficient = −0.1254, p < 0.001). We tested how MiSeq runs of 2 x 300 or 2 x 330-cycles affected sequence retention (Fig. S1). Although the 2 x 330-cycle run generated more raw data, it produced shorter Q30-trimmed read pairs than the 2 x 300-cycle run (Fig. S1A, p = 6.2 x 10^−6^), with the decrease primarily in read 2 (Fig. S1B, p = 3.2 x 10^−6^). All following analyses were produced from the 2 x 300-cycle sequencing data.

**Figure 2.**
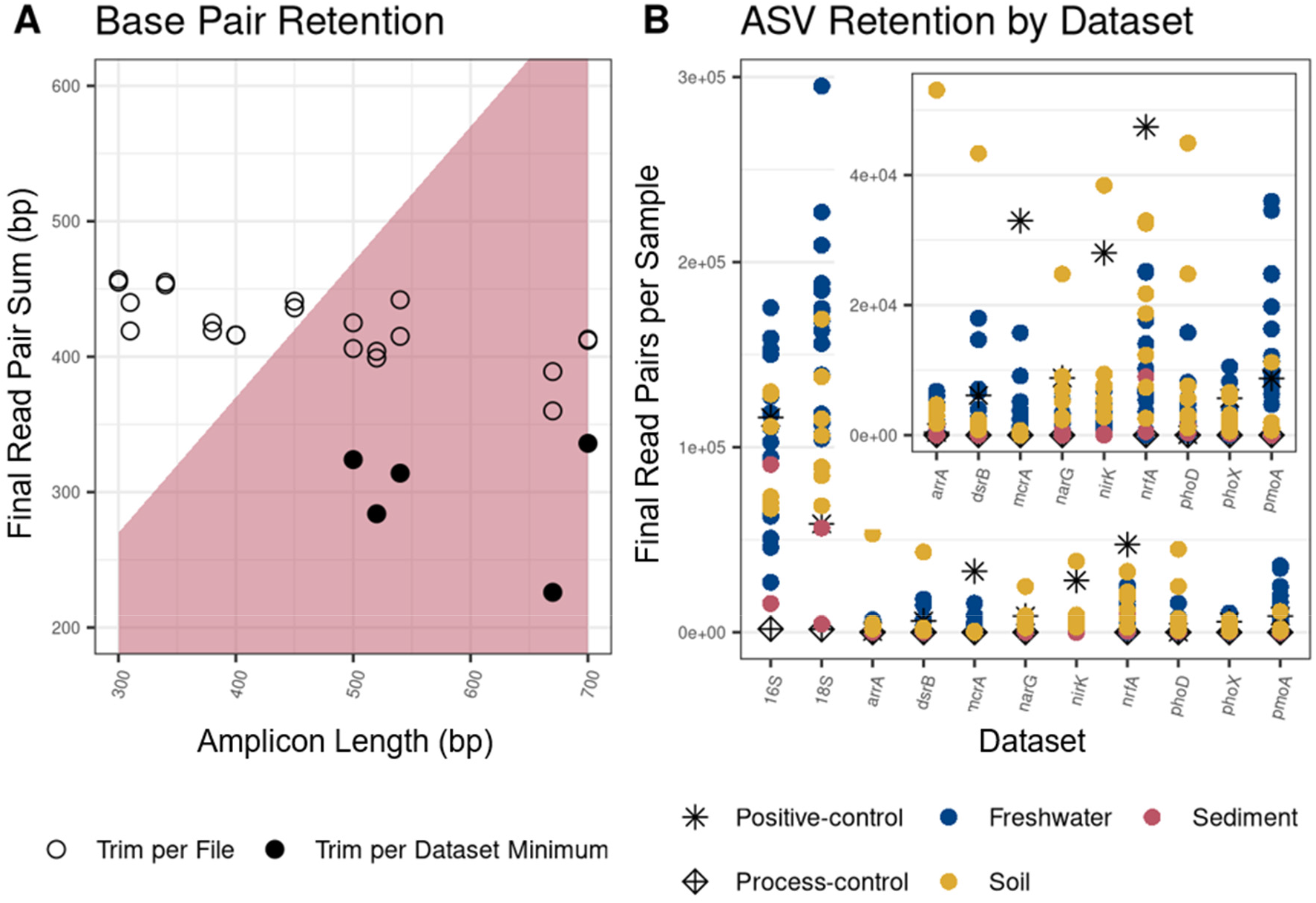
Data retention by dataset. A) Sequence lengths obtained for each amplicon plotted against expected amplicon length. Red shading denotes read pairs that must be concatenated instead of merged due to insufficient bp retention. Final read pair sum (bp) is the sum of bases retained per read pair of a given orientation after each file was trimmed to Q30 with the Trim per File or Trim per Dataset Minimum methods described in the text. Trim per Dataset Minimum is only shown for concatenated datasets. B) Sequence data retention per sample as the sum of final read pairs plotted for each dataset. Sample type is indicated by symbol and color as indicated in the legend. The panel inset expands the y-axis to better show differences for functional gene datasets.

The positive control contained a staggered genomic mix and individual genomes containing functional genes of interest with normalized copy numbers to assess amplicon and sequencing sensitivity (Table S3). 16S rRNA sequences were observed for all members of the positive control except *Enterococcus faecalis,* which had the lowest relative abundance of 0.03%. *Pseudomonas fluorescens* was also present 0.03% relative abundance and was detected. Functional gene detection in the positive control was initially assessed by the presence of an appropriate size amplicon. Because amplicons in the *phoD_X* fragment analysis were outside the predicted *phoD* range, the eight most abundant ASVs in the *phoD* sequence dataset were used to search the NCBI non-redundant nucleotide database by BLASTN. The top hits were known *phoD* sequences with the best match having a maximum nucleotide identity of 70% and an expect value of 2×10^−50^.

### Sample and Amplicon Factors Affecting Sequence Retention

We examined factors affecting sequence retention by CAMASE (Figure 2B). CAMASE sequences higher concentrations of *phoD_X* and *narG* amplicons to offset amplicon pooling and large size, respectively. Even so, the shortest amplicons, *nrfA* and 18S rRNA, produced the largest number of final read pairs indicating preferential sequencing of short amplicons. Sample type influenced sequencing yield (Table 2). All soil samples had optimal template, while half the freshwater and both sediment samples were sub-optimal. The *nirK* dataset had different sequence retention comparing freshwater and soil samples (p=0.016) while *narG* (p=0.030) and *nirK* (p=0.010) had significantly increased retention in optimal vs. sub-optimal template groups. Increased retention for the optimal template was observed for *arrA*, *dsrB*, *nrfA*, *phoD*, and *phoX*, albeit not significantly (p > 0.05).

### Data Overview and Trends

CAMASE produces analyses for each dataset that can be organized by taxonomic rank or OTU cluster similarity (Fig. 3). The data presented here are for 16S rRNA as the most common amplicon dataset, *dsrB* as a dataset where reads 1 and 2 were merged by overlap, and *nirK* where reads 1 and 2 were concatenated.

**Figure 3.**
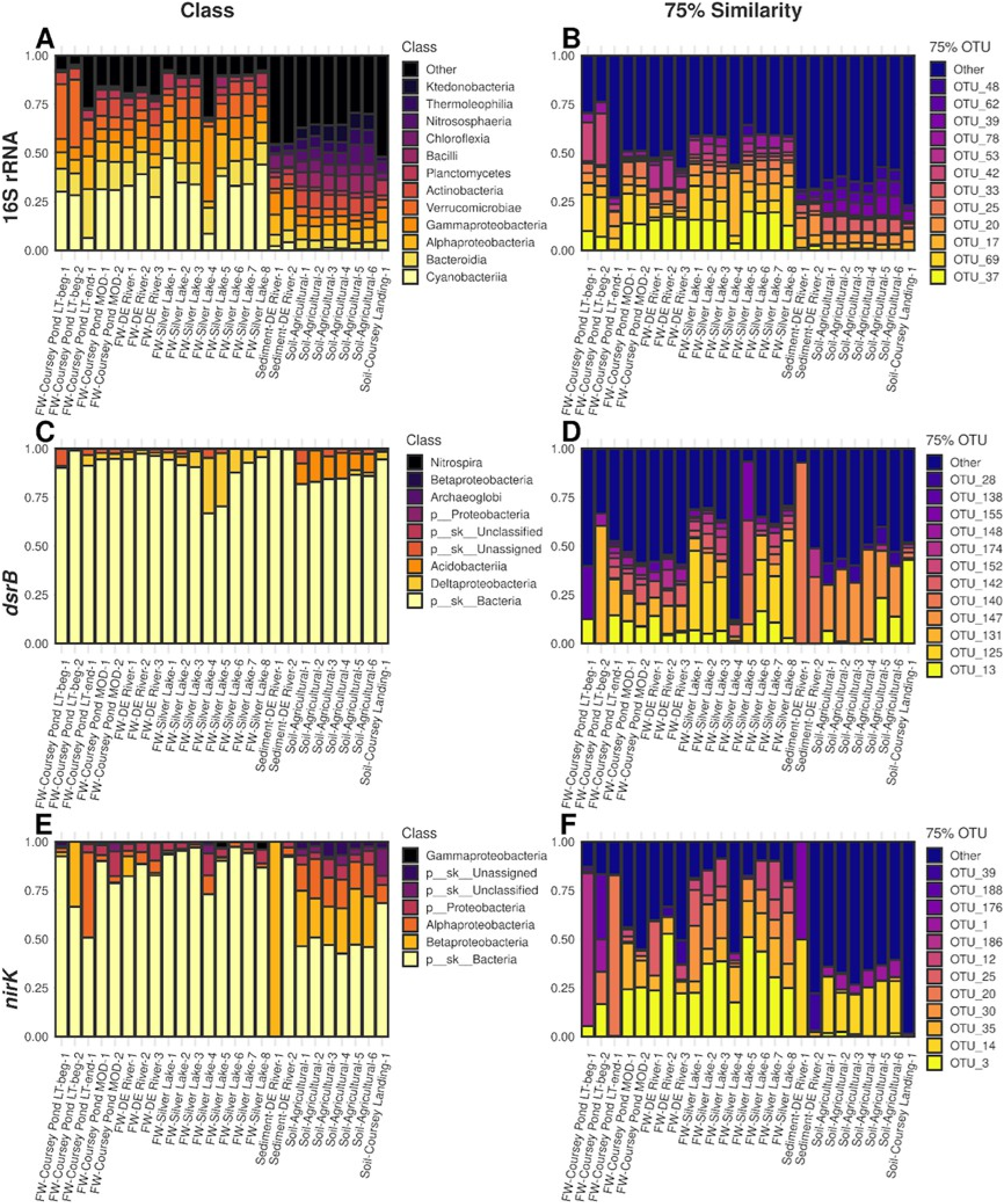
Relative abundance for selected datasets by class (A, C, E)or 75% similarity OTUs (B,D,F). For simplicity, the results of 16S rRNA (A, B), *dsrB* (C, D), and *nirK* (E, F) are displayed. Plots for all datasets are in Supplemental Material. For panels A,C, and E, the assigned taxonomy for any sequence was propagated from the lowest level assigned by the classifier.

Clustering by Class in 16S rRNA (Fig. 3A) provides a familiar summary of sequence classification and indicated Cyanobacteria are the dominant Class in freshwater samples, while soil samples do not have a clearly dominant taxon. For *dsrB* (Fig. 3C) and *nirK* (Fig. 3E), the dominant Class is p sk Bacteria. This rank is automatically generated by CAMASE and indicates that these sequences are from Bacteria, but could not be confidently assigned to a lower taxonomic rank.

Patterns of OTU relative abundance at 75% similarity show greater numbers of OTUs relative to Class for all genes. For 16S rRNA (Fig. 3B), we can infer that OTU_37 corresponds to Class Cyanobacteria while OTU_33 corresponds to Class Verrucomicrobia based on the patterns observed freshwaters while OTU_17 and OTU_20 correspond to Classes Alpha- and Gammaproteobacteria based on soil samples. OTU clustering provides more nuance for functional genes. For *dsrB* (Fig. 3D), OTU_125 and OTU_131 are prevalent in freshwater while OTU_147 is characteristic for soils, while OTU_140 is associated with the two sediment samples. For *nirK* (Fig. 3F), OTU_3 is characteristic for freshwaters while OTU_14 is characteristic for soils. Sediment sample DE River-1 *nirK* sequences are dominated by Class Betaproteobacteria *nirK* (Fig. 3E), while OTU clustering indicates this can be split into two separate *nirK* sequence types represented by OTU_14 and OTU_1. OTU_1 is widely present, but was only dominant in this sediment sample. Plots for all remaining datasets at the Class, ASV, and OTU at 95%, 85%, and 75% similarity cutoffs along with code used to generate them are provided (Supplemental File 2). Detailed markdowns available at the CAMASE git instance can be modified to produce plots for taxonomic levels from Domain to Genus and any user specified similarity cutoff.

CAMASE employs compositional data treatment by performing Principal Component Analysis (PCA) using the Aitchison distance, which requires zero replacement. PCA analysis was performed on ASVs (Fig. 4A, D, G), OTUs at multiple similarity thresholds (75% shown, Fig. 4B, E, H), and multiple taxonomic ranks (Class shown, Fig. 4C, F, I). Plots for the remaining datatsets are provided in Supplemental Material (Supplemental File 2). With 16S rRNA, similar levels of variance (39.7-43.6%) were captured in the first two principal components regardless of how data were grouped. Functional genes tended to capture less variance with ASVs or 95% OTUs (29-36%), while grouping by Class captured more variation (56-80%). This reflects taxonomy assignment issues for functional genes that are discussed below. Functional genes showed different patterns of sample separation. For example, Delaware River samples were resolved along PC2 with *dsrB* 95% OTUs (Fig. 4E, inverted triangles), while all other sample types were tightly clustered. In contrast, agricultural soils were well separated along both PC1 and PC2 by *nirK* (Fig. 4H, gold stars), while all others clustered by sample type.

**Figure 4.**
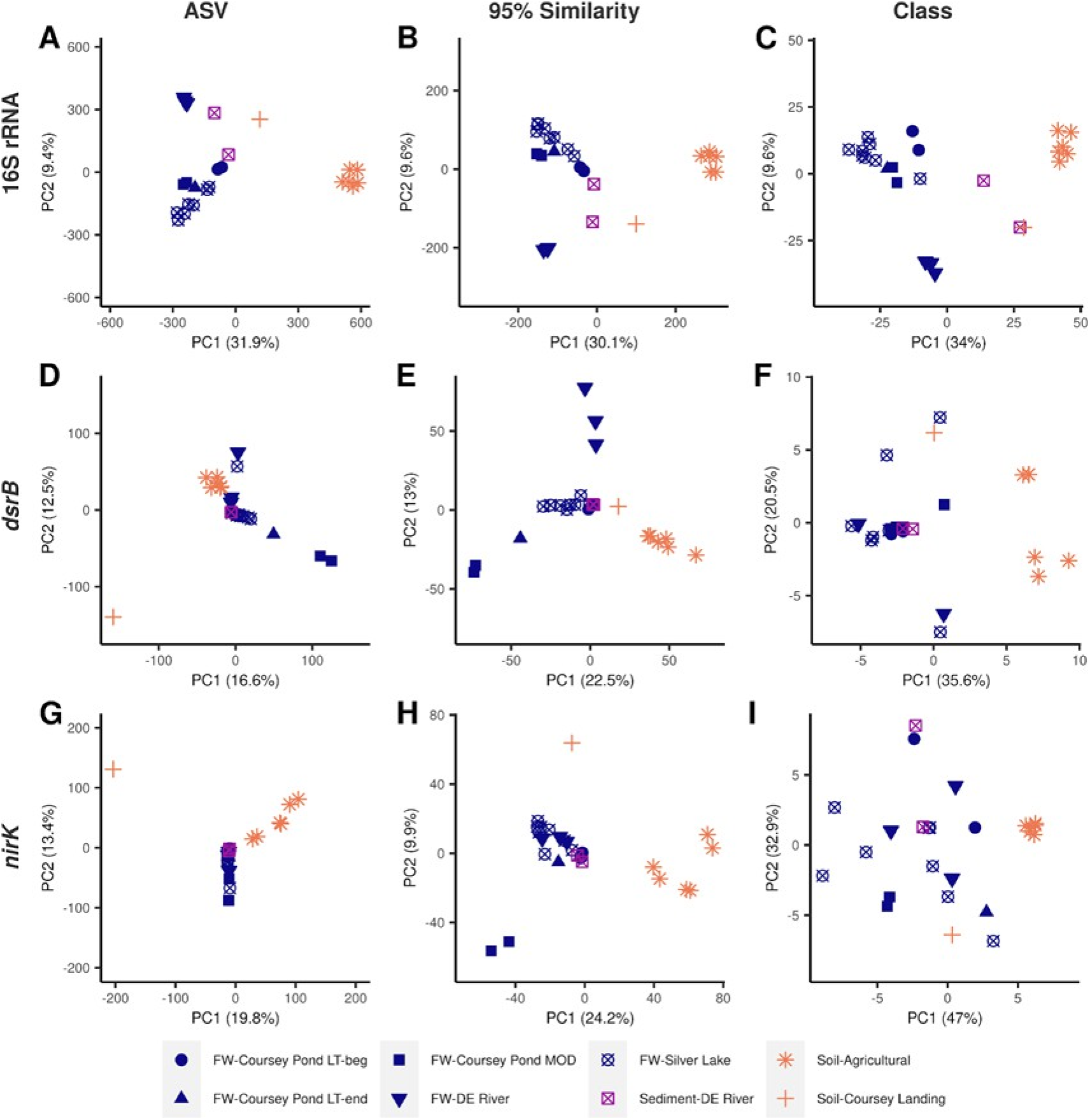
PCA for selected datasets clustered as ASVs (A, D, G), 95% OTUs (B, E, H), or Class (C, F, I). For simplicity, 16S rRNA (A, B, C), *dsrB* (D, E, F), and *nirK* (G, H, I) are displayed here with plots for all datasets available in Supplemental Information. Data in panels C, F, and I were calculated after collapsing to class with NA taxonomic assignments propagated from the lowest taxonomic level assigned by the classifier. The axes indicate the principal component followed by the percentage of variance explained.

Alpha diversity metrics were calculated with DivNet [14], which uses a parameter called perturbation for zero replacement with a default value of 0.05. Very sparse datasets, where log(number of zeros / sum of observations) > 0, produced very high diversity estimates with the default perturbation value at any OTU clustering threshold (Fig. 5). Therefore, we varied the perturbation down to 5×10^−5^. Shannon estimates converged to expected low values for samples with log(number of zeros / sum of observations) < 6 when the perturbation values was set to 1×10^−4^ (Fig. 5).

**Figure 5.**
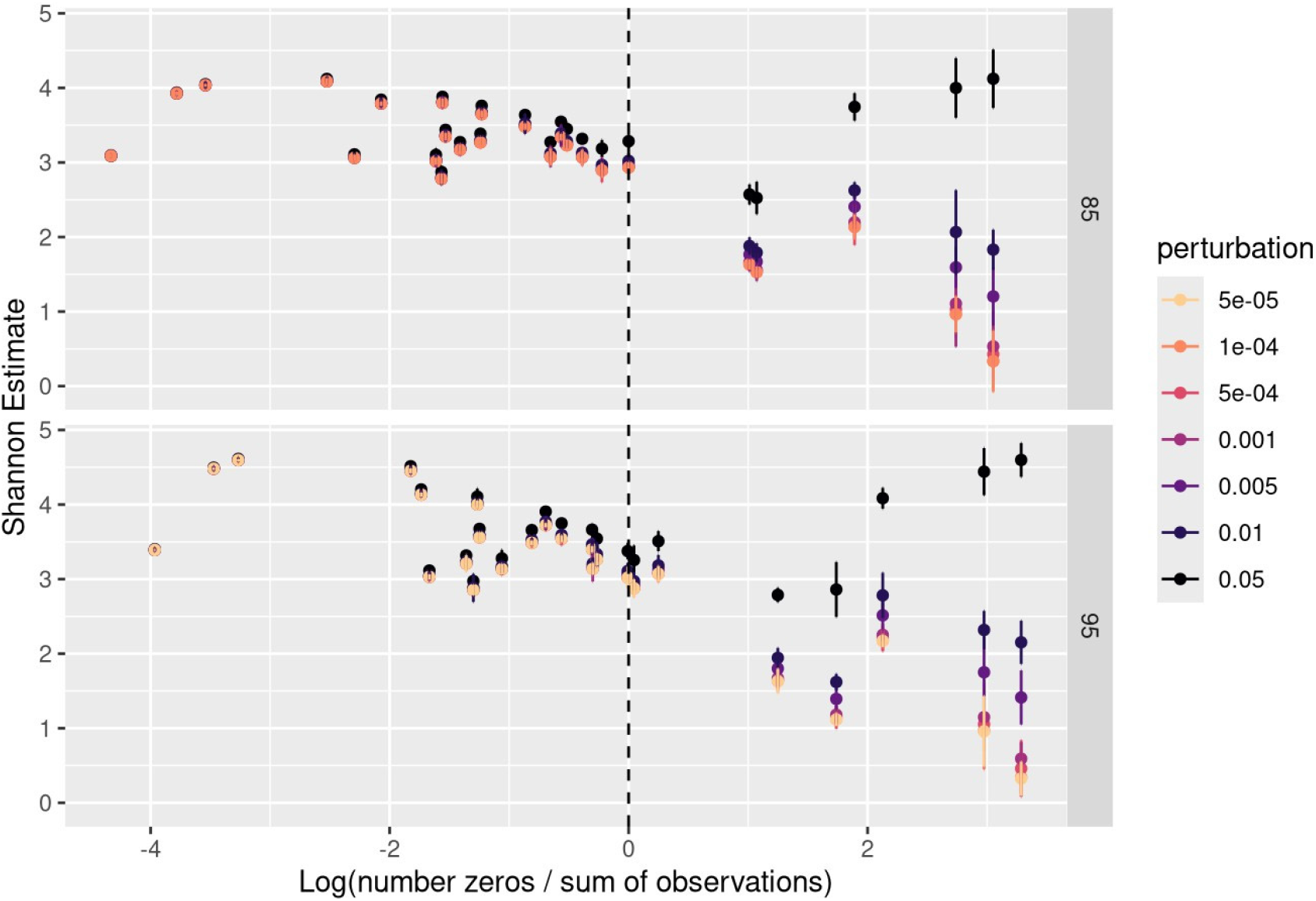
Effects of perturbation value selected on Shannon diversity estimate. The example shown is *dsrB* OTUs clustered at 85% or 95% similarity. Results for all datasets are found in Supplemental Information (Supplemental File 3).

Estimated Shannon diversity varied with sample type in all datasets (Fig. 6A). Soils typically had the highest estimates, followed by freshwaters and sediments, except for 16S rRNA, where and freshwaters and sediments were similar. Finer scale patterns could be distinguished; for example, the lowest diversity of freshwater samples were during Cyanobacterial blooms in Coursey Pond (FW-Coursey Pond-LT-beg/end). Shannon diversity for 16S rRNA and 18S rRNA were positively correlated (Fig. 6B), as were 16S rRNA and *nirK* (Fig. 6D), *nrfA*, *phoD*, or *phoX* (Supplemental File 3). 16S rRNA and *mcrA* estimates were negatively correlated, but not significantly (Supplemental File 3). The remaining six functional genes did not correlate with 16S rRNA diversity, as shown for *dsrB* (Fig. 6C).

**Figure 6.**
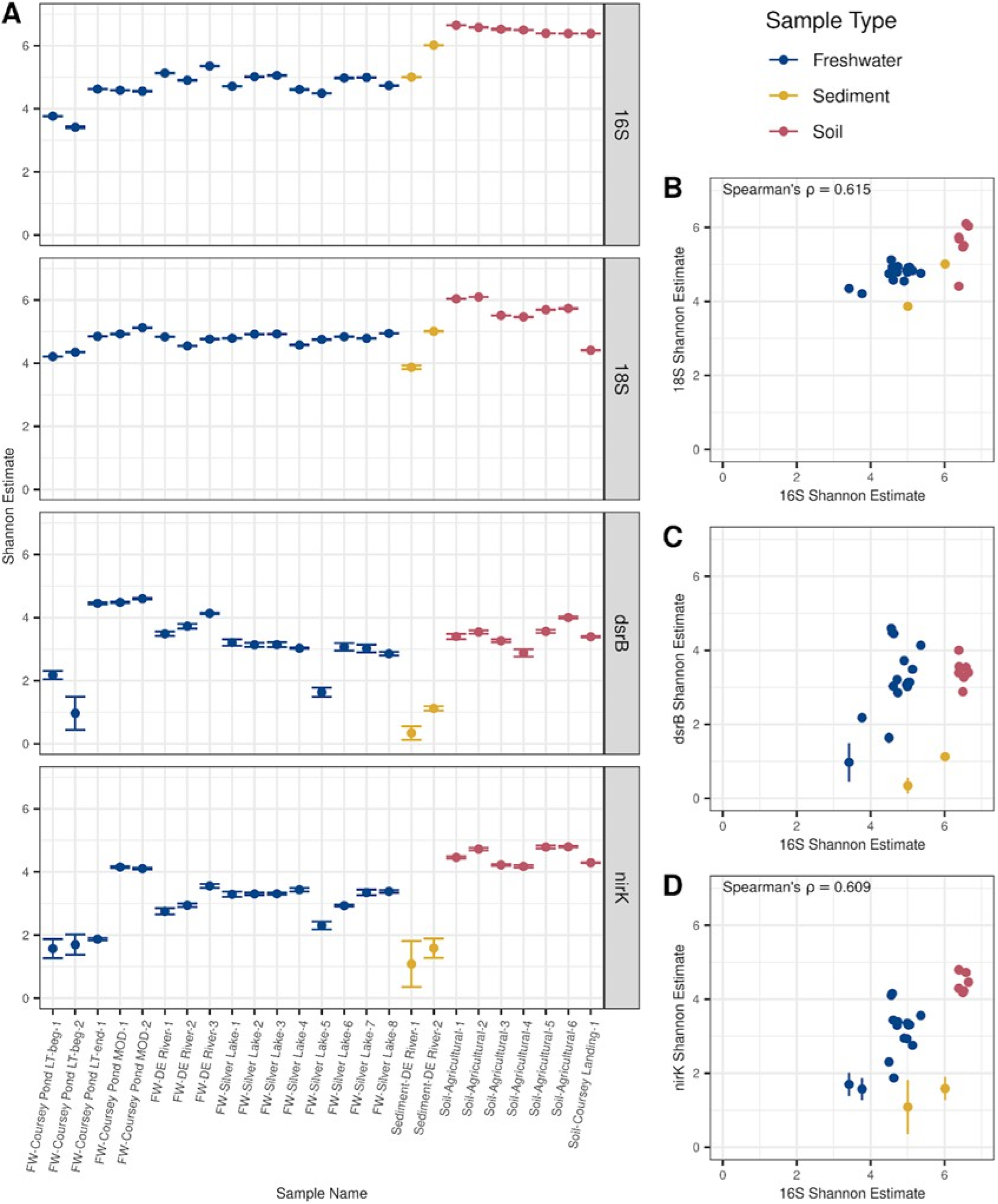
Diversity estimates across samples and correlation between datasets across all samples. A) DivNet estimated Shannon diversity and 95% confidence intervals (CIs)for each sample within the 16S rRNA, 18S rRNA, *dsrB*, and *nirK* datasets. Values were calculated with the sample name as the only covariate. Panels B, C, and D plot the 16S Shannon estimate with 95% CIs on the x-axis against the estimate for 18S (B), *dsrB* (C), or *nirK* (D) with 95% CIs along the y-axis, respectively. Spearman’s rank correlation of the mean estimates with Bonferroni correction, ɑ = 0.05, p-value 0.0134 (18S), 1 (*dsrB*), and 0.0153 (*nirK*); Spearman’s ⍴ is displayed for statistically significant correlations.

### CAMASE is Reproducible

Five DNA samples, three soils and two waters, were processed twice starting at gene-specific PCR and the Aitchison distances between 95% OTUs of replicates were evaluated. The Mantel r was > 0.99 (p = 0.001) for 16S rRNA, 18S rRNA, and *phoD* indicating that a given OTUs was observed at nearly the same abundance in each replicate. The Mantel r was > 0.92 for all functional marker genes (p = 0.001). Only *arrA* and *dsrB* had moderate Mantel r values, 0.539 (p = 0.19) and 0.873 (p = 0.001), respectively. The test power is less than 0.8 for all comparisons and was most likely limited by the number of samples replicated [35]. Thus, we also qualitatively evaluated by inspecting PCA plots (Fig. 7). In all datasets, technical replicates were located closer to one another than to other samples of the same environmental type.

**Figure 7.**
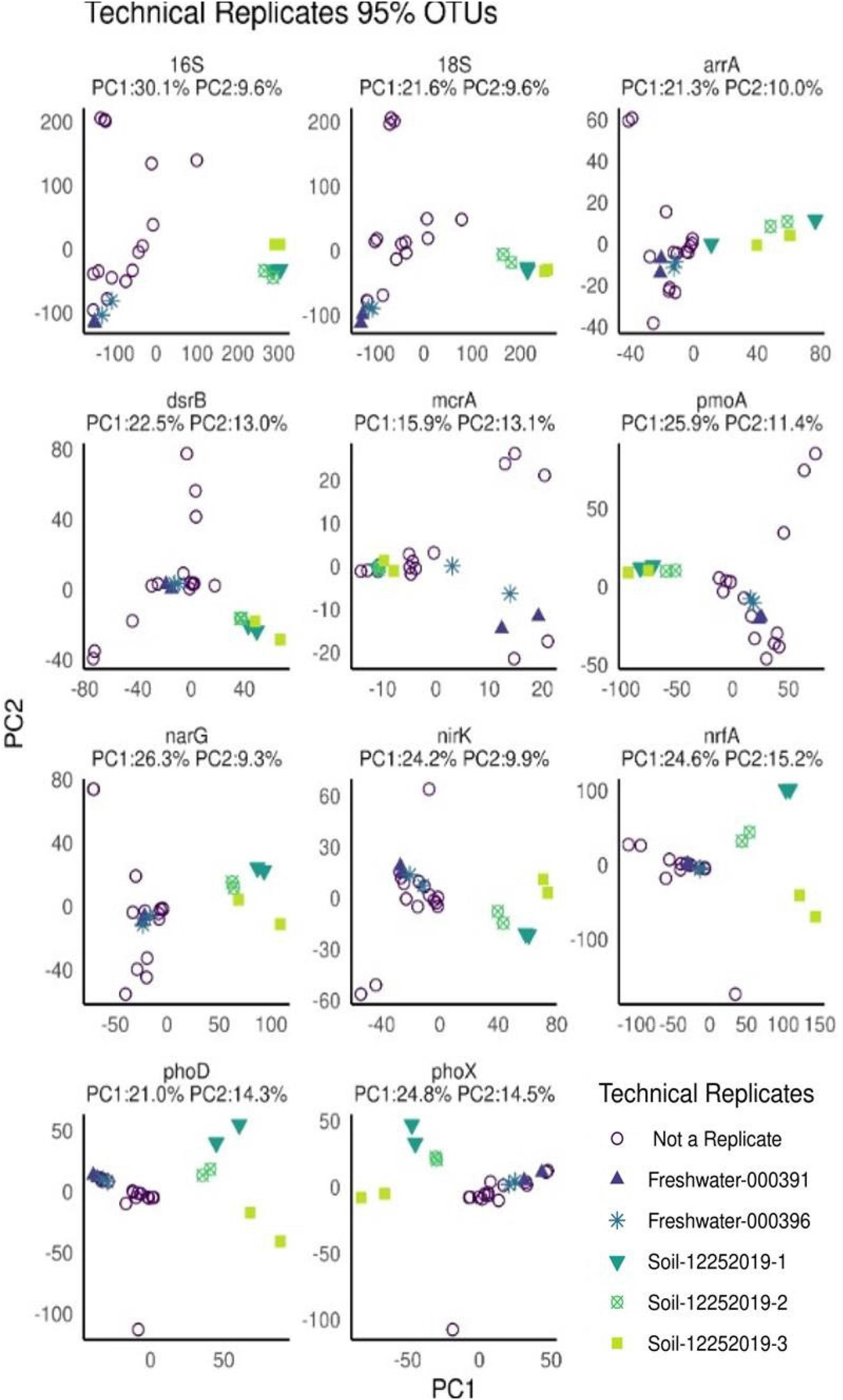
PCA of technical replicates clustered at 95% similarity. Environmental samples with technical replication are distinguished by color and shape. All samples which did not have technical replicates are shown as open circles. The percentage of variance explained by each principal component is denoted at the top of each graph.

## DISCUSSION

The CAMASE workflow enables the parallel production, sequencing, and analysis of eleven phylogenetic and functional gene amplicons from individual samples collected across diverse sample types. This study was designed as a proof-of-concept with diverse samples rather than a powered ecological comparison with statistical tests mainly interpreted as performance diagnostics. Outputs from reproducibility testing identified workflow aspects to consider when using CAMASE. With this context, CAMASE produced highly similar profiles for nine out of eleven amplicons in technical replicates and worked well with 1 ng or less DNA template per gene-specific PCR reaction. Finally, CAMASE was designed to be intercomparable with existing data by using published primer sets. For example, 16S and 18S rRNA amplicons were designed to enable inter-study comparisons with the Earth Microbiome Project [8].

CAMASE standardizes sample collection and DNA extraction methods regardless of sample type. It integrates robotic liquid handling for PCR amplification and pooling. Both choices were made to improve consistency. With the two-step barcoding approach used here, it has been shown that sample preparer can be the largest source of between-replicate variation [12]. The workflow can be translated to other models of liquid handling robots that may be available in individual labs or core facilities. It can also be carried out with multi-channel pipettors for laboratories without access to automated liquid handling, albeit with a chance for increased variability.

Taxonomic (16S and 18S rRNA) and functional genes (all other datasets) had differing sequence retention (Table 1, Fig. 2B). CAMASE collects larger amounts of taxonomic gene data from every sample by including 16S and 18S rRNA amplicons at twice the concentration of most other amplicons to account for the fact that these genes are universal and deeper sequencing is required to fully sample taxonomic gene diversity than functional gene diversity. This is because a given functional gene is not expected to be present in every cell in a sample or evenly distributed across all sample types. We also observed amplicon size effects on sequence retention where the two smallest amplicons, *nrfA* and 18S rRNA, had higher numbers of final read pairs (Table 1, Fig. 2B). Future iterations of CAMASE could further explore and optimize for amplicon size.

Variability in sequence retention by sample type was expected because we challenged CAMASE with diverse sample types. Environmental factors are known to impact biogeochemical gene distribution, as well documented in soils [36]. The samples here contained two different soil types along with sediments and freshwaters from multiple sites across Delaware. Even so, CAMASE produced interpretable data from this broad collection. Expected trends were observed like the high relative abundance of Cyanobacteria in freshwater samples and increased diversity of soil samples relative to sediments and waters. Users adopting CAMASE for narrower sample types can optimize pooling, sequencing depth, and sequence filtering thresholds for their uses.

Sequencing and sequence data processing affected both data quantity and quality produced by CAMASE. We observed an inherent trade-off between quantity of bases and quality, and CAMASE favors quality. We strongly recommend 2 x 300-cycle sequencing and data processing with trimming per file. With this, amplicons < 400 nt could be merged by overlapping sequence reads, while those > 500 nt required concatenation. Merging of overlapping sequences provides additional quality control, as recommended by the developers of DADA2 [20]. Amplicons between 400 and 500 nt must be assessed on a run-by-run basis. The data presented show that read 2 quality drives overall sequence retention and will determine the final decision to merge vs. concatenate in the 400-500 bp window.

Shannon diversity estimates for longer amplicons that are concatenated should be evaluated cautiously. Because amplicon orientation is not fixed in the CAMASE workflow, the F and R gene-specific primers may be in either read 1 or read 2, resulting in distinct merged datasets called 1F2R and 1R2F. CAMASE currently merges all ASVs for 1F2R and 1R2F into a single observational count table. If quality trimming and merging are not identical for the same amplicon sequence in different read orientations, this will result in two ASVs in place of one. Future iterations of CAMASE could test using two observational count tables covering both orientations as pseudo-replicates.

CAMASE produced highly similar OTU profiles for technical replicates. The Mantel r values were > 0.873 for all datasets, except *arrA*. However, the power was less than 0.8 for all tests; a power between 0.8 and 0.9 is recommended to balance type I and type II errors [35]. The lower power observed here indicates a low chance that high r-values are false positives and that most amplicons are reproducible in this proof-of-concept study. We interpret the *arrA* data as reflecting the diverse sample type analyzed and low replication level used rather than an inherent flaw in the *arrA* primer set. This is supported by PCA plots showing that technical replicates cluster tightly in all datasets, including *arrA*. Users are encouraged to critically evaluate technical reproducibility when adopting CAMASE for their own use.

Computational tools for amplicon sequencing analysis have been designed and optimized for rRNA and their utility for functional genes is not yet established. Functional gene datasets posed challenges. Most notable is the dearth of taxonomic assignments for functional gene amplicons in reference sequence databases. This means that CAMASE assigns 50 - 90% of functional gene amplicon OTUs to the Domain Bacteria with no classification at any lower taxonomic rank. Our best advice is to examine functional gene OTU abundances across samples to identify trends and characteristic functional gene OTUs for specific samples. CAMASE captures the centroid sequence for every ASV and OTU that can be used for more detailed taxonomic analyses. Hidden Markov Models for sequence classification may also provide more sensitive and specific taxonomic assignment at lower sequence identities than current classifiers [37, 38].

CAMASE uses DivNet for diversity metric estimation with confidence intervals [14]. We made two key modifications to default parameters. First, the perturbation value was adjusted from 0.05 to 0.0001 based on a sensitivity test with the CAMASE outputs. This reflects that zero values for multiple OTUs across the diverse sample types in this proof-of-concept are plausible and avoids assigning excessive weight to these features. The second was developing a wrapper function to pick the base OTU used for calculations. DivNet expects to find one or more OTUs that are present in every sample in the data provided, but this is also not a reasonable expectation for the diverse sample types analyzed here. For hypothesis driven ecological studies using CAMASE, the same assumptions are likely not valid and should be explored by users using methods outlined here.

With these adjustments Shannon diversity metrics had two patterns of note. First, soils generally had higher Shannon diversity relative to sediments and freshwaters. This seems intuitive given the high spatial heterogeneity and more stable physical structure within soils relative to waters and surficial sediments [39]. Second, the Shannon diversity for *narG*, *nirK*, *nrfA*, *phoD*, and *phoX* were significantly positively correlated with 16S rRNA diversity. These genes encode enzymes for N and P cycling. This may reflect a broad phylogenetic distribution of these genes making them reliable reporters of overall community shifts. Weak correlations for other genes underscore the value of directly measuring functional-gene diversity rather than assuming it scales with taxonomic diversity.

CAMASE is adaptable to different research interests and is sensitive. Any primer set can be substituted by synthesizing it with the appropriate 5’-end extension on both forward and reverse primers. We recommend selecting primers producing amplicons < 450bp—including barcodes—resulting in 350-400 bp after trimming for the benefit of read pair merging. The data here show that CAMASE works with low sample inputs and the control samples were intended to monitor amplification and processing performance, not contamination. We encourage users to enable contaminant removal by including at least three controls per sequence library, two process controls and the reagent negative, to enable contaminant-aware filtering via software packages like Decontam [40].

CAMASE was designed as a cost-effective middle ground between metagenomics and single amplicon studies. CAMASE as employed here produced 11 amplicon datasets for 25 samples at a cost of $5,600 when the sequencing was conducted in 2019. This is less than $225 per sample and less than $21 per amplicon within a sample. Local pricing for consumables, reagents, and sequencing costs will strongly influence the cost. Primer synthesis for the two-step barcoding approach is the biggest initial cost with sequencing runs being the largest recurring cost. Sequencing technology continues to evolve and with costs decreasing and throughput increasing [41]. Transitioning CAMASE to the AVITI sequencing platform, which offers higher throughput and equal or better per-base accuracy than MiSeq [42], is being investigated.

CAMASE’s computational workflow uses a single discrete explanatory variable: sample type. This can be modified for other explanatory variables by editing sample metadata and CAMASE scripts from the git instance. Access to computational power can limit CAMASE primarily due to DECIPHER alignment/OTU clustering and DivNet diversity calculations. A moderately powerful linux workstation (6 dual-threaded cores at 3.6 GHz, 64 GB RAM) was able to run DivNet on the 85% OTU cluster while a single high-performance computing cluster node (32 dual-threaded cores at 2.8 GHz, 512 GB RAM) enabled DivNet calculations on 95% similarity OTU clusters. The recent heuristics released for clustering with DECIPHER should reduce this time by significantly [43]. A RUST implementation of DivNet with increased CPU and memory efficiency for alpha diversity calculations [44] should also improve performance. Ultimately, analyzing sample types that are less diverse than those used in this proof-of-concept should produce fewer OTUs which is the main driver of computation time.

## SUMMARY

We provide a proof-of-concept for CAMASE as a workflow for microbial community diversity and functional potential by compositional analysis of multiple amplicon sequence datasets. CAMASE encompasses sample collection, preservation, extraction, amplification, multiplexing, sequencing, and computation. Water, soil, and sediment samples produced technically reproducible results with low DNA template input. We feel the CAMASE approach is a good starting point to identify samples worth investing in deep metagenomic sequencing to develop genome-resolved functional inferences. The workflow available at the CAMASE git instance includes standard operating procedures (SOPs), calculation templates, robotic programs, and bioinformatic scripts. The computational analysis was designed to be complete while remaining accessible to non-bioinformaticians. This could make CAMASE particularly attractive for courses that want to expose students to both the laboratory and computational aspects of molecular microbial ecology or research laboratories just venturing into the field.

## Supporting information

Sample metadata

Markdown for relative abundance and PCA plots

Markdown for diversity estimates

## DATA AVAILABILITY

Raw sequencing reads have been deposited at the NCBI Sequence Read Archive and can be accessed under BioProject Accession PRJNA1473962. Markdown and metadata files used to produce the figures in this manuscript from the sequence reads from the 2 x 300 bp reads as well as detailed instructions and metadata templates are available at https://hansonlabgit.dbi.udel.edu/aprange/CAMASE.

## ACKNOWLEDGMENTS

The authors would like to thank the University of Delaware’s DNA Sequencing and Genotyping core facility, particularly Mark Shaw, for performing Illumina MiSeq library preparation and sequencing runs. The authors thank Miranda Marini, Alena Brown and Alwayne Allen for testing initial versions of sample collection and DNA extraction protocols. Cas Derieux, Ray Loughran, and Aerin Rost-Nasshan are thanked for providing constructive feedback on a manuscript draft. This publication was made possible by the National Science Foundation EPSCoR Grant No. 1757353 and the State of Delaware. Support from the University of Delaware Bioinformatics Data Science Core Facility (RRID:SCR_017696) including use of the BIOMIX and BioStore computational resources was made possible through funding from National Institutes of Health Shared Instrumentation Grant (NIH S10OD028725), the State of Delaware, and the Delaware Biotechnology Institute. Any opinions, findings, and conclusions or recommendations expressed in this material are those of the author(s) and do not necessarily reflect the views of the National Science Foundation, National Institutes of Health, or State of Delaware.

## Author Roles

AB: Data curation, formal analysis, investigation, visualization, writing – original draft and review & editing; RM: Software, writing – review & editing; CH: Software, writing – review & editing; TH: Conceptualization, funding acquisition, project administration, writing – original draft and review & editing.

## Notes

### Competing Interest Statement

The authors have declared no competing interest.

