## Supplemental text, tables, and figures for "A Sample to Results Workflow for Compositional Analysis of Multiplexed Amplicon Sequencing Experiments"

**Brief Descriptions of Supplemental Files:**

**Supplemental File 1:** A .csv file containing metadata for controls and environmental samples analyzed in this manuscript.

**Supplemental File 2:** A .html file containing markdown of code and both relative abundance and PCA plots for datasets not presented in the main text.

**Supplemental File 3:** A .html file containing markdown of code and plots for Shannon diversity estimates of datasets not contained in the main text.

**Table S1.** Gene-specific primer sets used in the first PCR step.

| Dataset | Primer Name | Sequence (5'-3') | Reference |
|---|---|---|---|
| ***Forward*** | | | |
| *pmoA* | A189 | GGNGACTGGGACTTCTGG | [1] |
| *arrA* | arrA-CVF1 | CACAGCGCCATCTGCGCCGA | [2] |
| *dsrB* | DSR1762Fmix_modified | equimolar parts DSR1762F1-5,7-9 (Table S2) | [3, 4] |
| *gcd* | gcd-FW | CGGCGTCATCCGGGSITIYRAYRT | [5] |
| *mcrA* | MCRf | TAYGAYCARATHTGGYT | [6] |
| *narG* | narG1960f | TAYGTSGGSCARGARAA | [7] |
| *nirK* | nirKC1F | ATGGCGCCATCATGGTNYTNCC | [8] |
| *nrfA* | nrfAF2aw | CARTGYCAYGTBGARTA | [9] |
| *phoD_X* | phoD_XFmix | equimolar parts phoD-F733 and phoX-F455 (Table S2) | [10, 11] |
| *phoN* | phoN-FW | GGAAGAACGGCTCCTACCCIWSNGGNCA | [5] |
| 18S rRNA | S-*-Univ-1391-a-S-16 (1391F) | GTACACACCGCCCGTC | [12] |
| 16S rRNA | S-D-Bact-0515-b-S-19 | GTGYCAGCMGCCGCGGTAA | [13] |
| ***Reverse*** | | | |
| *pmoA* | mb661 | CCGGMGCAACGTCYTTACC | [14] |
| *arrA* | arrA-CVR1 | CCGACGAACTCCYTGYTCCA | [2] |
| *dsrB* | DSR4Rmix_modified | equimolar parts DSR4R a-g and DSR698R (Table S2) | [3, 4] |
| *gcd* | gcd-RW | GGGCATGTCCATGTCCCAIADRTCRTG | [5] |
| *mcrA* | MCRr | ACRTTCATNGCRTARTT | [6] |
| *narG* | narG2650r | TTYTCRTACCABGTBGCACCABGTBGC | [7] |
| *nirK* | nirKC1R | TCGAAGGCCTCGATNARRTTRTG | [8] |
| *nrfA* | nrfA-R1 | TWNGGCATRTGRCARTC | [15] |
| *phoD_X* | phoD_XRmix | equimolar parts phoD-R1083 and phoX-R1076 (Table S2) | [10, 11] |
| *phoN* | phoN-RW | CACGTCGGACTGCCAGTGIDMIYYRCA | [5] |
| 18S rRNA | EukBr | TGATCCTTCTGCAGGTTCACCTAC | [16] |
| 16S rRNA | S-D-Bact-0806-b-A-20 | GGACTACNVGGGTWTCTAAT | [17] |

**Table S2.** Summary of primer mixtures for *dsrB* and *phoD_X* amplification.

| Dataset | Primer Name | Sequence (5'-3') | Reference |
|---|---|---|---|
| ***Forward*** | | | |
| *dsrB* | DSR1762F1 | CAYACCCAGGGNTGG | [3, 4] |
| | DSR1762F4 | CACACDCAGGGNTGG | |
| | DSR1762F2 | CAYACBCAAGGNTGG | |
| | DSR1762F3 | CATACDCAGGGHTGG | |
| | DSR1762F5 | CATACHCAGGGNTAY | |
| | DSR1762F7 | CACACBCAGGGMTAC | |
| | DSR1762F8 | CACACHCAGGGCTAT | |
| | DSR1762F9 | CACACCCAGGGWTT | |
| *phoD_X* | phoD-F733 | TGGGAYGATCAYGARGT | [10] |
| | phoX-F455 | CAGTTCGGBTWCAACAACGA | [11] |
| ***Reverse*** | | | |
| *dsrB* | DSR4Ra | GTGTAACAGTTTCCACA | [3, 4] |
| | DSR4Rb | GTGTAACAGTTACCGCA | |
| | DSR4Rc | GTGTAGCAGTTKCCGCA | |
| | DSR4Rd | GTGTAGCAGTTACCACA | |
| | DSR4Re | GTGTAACAGTTACCACA | |
| | DSR4Rf | GTATAGCARTTGCCGCA | |
| | DSR4Rg | GTGAAGCAGTTGCCGCA | |
| | DSR698R | GTGTARCAGTTRCCRCA | |
| *phoD_X* | phoD-R1083 | CTGSGCSAKSACRTTCCA | [10] |
| | phoX-R1076 | CGGCCCAGSGCRGTGYGYTT | [11] |

**Table S3.** Positive control DNA mixture components.

| Genome Mix | Target | Item number | batch | DNA (ng) | PCR copy #[1] |
|---|---|---|---|---|---|
| 10 Strain Mix | 16S rRNA | MSA-3002 | 70007470 | 260.15 | 400-400,000 |
| *Aspergillus fumigatus* | 18S rRNA | 1022D-2 | 70018097 | 0.011 | 800 |
| *Shewanella putrefaciens* | *arrA* | BAA-543-D | 58660978 | 0.031 | 380 |
| *Desulfovibrio vulgaris* | *dsrB* | 29579D-5 | 59679194 | 0.038 | 380 |
| *Escherichia coli* | *gcd, narG* | In MSA-3002 | 70007470 | 1.12 | 13,000 |
| *Methanosarcina acetivorans* | *mcrA* | 35395D-5 | 70008173 | 0.038 | 380 |
| *Alcaligenes faecalis* | *nirK* | 8750D-5 | 61646543 | 0.027 | 380 |
| *Wolinella succinogenes* | *nrfA* | 29543D-5 | 7410328 | 0.014 | 380 |
| *Bacillus subtilis* | *phoD* | ATCC 6051 | 70025850 | 0.028 | 380 |
| *Salmonella enterica* | *phoN* | 700720D-5 | 70039794 | 0.033 | 380 |
| *Methylococcus capsulatus* | *pmoA* | 33009D-5 | 70011371 | 0.022 | 380 |

1 – approximate number of copies of target gene present in gene-specific PCR

**Table S4.** Detailed PCR reaction conditions.

| **First Step PCR: gene specific** | **Second Step PCR: barcoding** |
|---|---|
| BioRad 384-well Thermal cycler<br>Method: Calc<br>Lid: 95°C<br>Volume: 20 uL<br>Total cycles: 30<br><br>Step Temp (°C) Time (min)<br><br>1 95 2:30<br>2 95 0:30<br>3 60 0:30<br>4 72 1:00<br>5 Goto step 2, 29x<br>6 72 5:00<br>7 4 ∞ | BioRad 384-well Thermal cycler Method: Calc<br>Lid: 95°C<br>Volume: 20 uL<br>Total cycles: 5<br><br>Step Temp (°C) Time (min)<br>1 95 2:30<br>2 95 0:30<br>3 60 0:30<br>4 72 1:00<br>5 Goto step 2, 4x<br>6 72 5:00<br>7 4 ∞ |

**Table S5.** Summary of fragment analysis results of gene-specific PCR reaction.

| | Size (bp)[1] | | |
|---|---|---|---|
| **Dataset** | **Exp** | **Obs** | **Comments[2]** |
| 16S | 320 | 327 | |
| 18S | 290 | 197 | |
| *arrA* | 360 | 389 | Major peak at 1,000 bp |
| *dsrB* | 430 | 408 | Minor additonal peaks >600 bp |
| *gcd* | 330 | ND | |
| *mcrA* | 520 | 565 | |
| *narG* | 680 | >650 | Major peak at 475 bp, multiple minor peaks >650 |
| *nirK* | 480 | 523 | |
| *nrfA* | 280 | 283 | |
| *phoD_X* | 380 & 650 | 645 | Minor peaks at 212 & 1089 bp |
| *phoN* | 160 | ND | |
| *pmoA* | 500 | 538 | Minor additional peaks |

1 - Amplicon size: exp = expected, obs = observed in fragment analysis, reported if +/- 10% of expected size, ND = not detected.

2 - Minor peak: <1/2 peak area of expected size, major: peak area ≥ peak area of expected size fragment

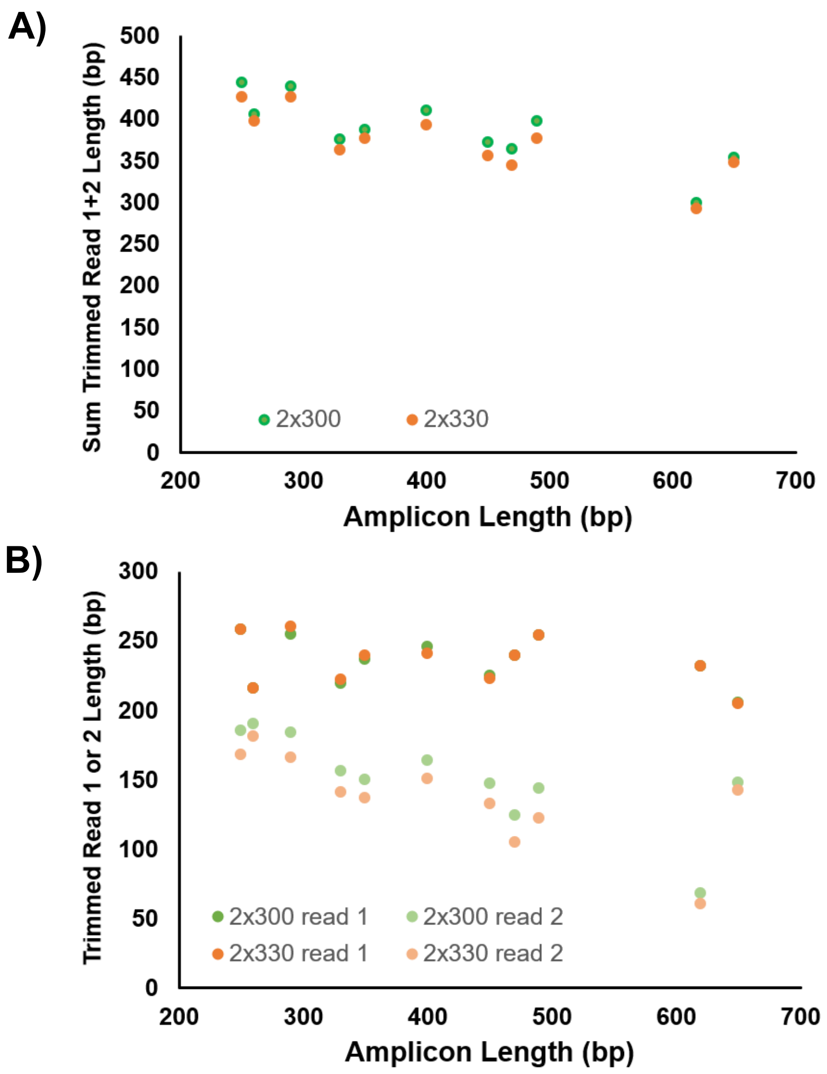


**Figure S1.** Read length comparisons between 2 x 300 and 2 x 330 cycle paired end MiSeq runs for amplicons in this study (Table S5). A) summed read 1 + 2 length; B) individual read 1 (dark points) or read 2 length (light points).
