## Supplementary material for "A Sample to Results Workflow for Compositional Analysis of Multiplexed Amplicon Sequencing Experiments": Markdown for relative abundance and PCA plots

Appendix A Figures All Samples Rel Abund & PCA


### Appendix A Figures All Samples Rel Abund & PCA

###### Alexa Bennett

#### 2024-12-22

###### Load Packages

```
library(magrittr)
library(biplotr)
library(tidyverse)
```

```
## ── Attaching core tidyverse packages ──────────────────────── tidyverse 2.0.0 ──
## ✔ dplyr     1.1.4     ✔ readr     2.1.5
## ✔ forcats   1.0.0     ✔ stringr   1.5.1
## ✔ ggplot2   3.5.1     ✔ tibble    3.2.1
## ✔ lubridate 1.9.3     ✔ tidyr     1.3.1
## ✔ purrr     1.0.2     
## ── Conflicts ────────────────────────────────────────── tidyverse_conflicts() ──
## ✖ tidyr::extract()   masks magrittr::extract()
## ✖ dplyr::filter()    masks stats::filter()
## ✖ dplyr::lag()       masks stats::lag()
## ✖ purrr::set_names() masks magrittr::set_names()
## ℹ Use the conflicted package (<http://conflicted.r-lib.org/>) to force all conflicts to become errors
```

```
library(featuretable) # Keep after tidyverse
```

```
## 
## Attaching package: 'featuretable'
## 
## The following object is masked from 'package:dplyr':
## 
##     collapse
## 
## The following objects are masked from 'package:purrr':
## 
##     keep, map
## 
## The following object is masked from 'package:biplotr':
## 
##     pca_biplot
```

```
library(viridis)
```

```
## Loading required package: viridisLite
```

```
library(cowplot)
```

```
## 
## Attaching package: 'cowplot'
## 
## The following object is masked from 'package:lubridate':
## 
##     stamp
```

###### Make some functions to clean-up chunks

```
make_unique <- function(x){
 if (anyDuplicated(x)){
   duplicates <- split(x, x)
   uniques <- names(duplicates)
   for (u in uniques){
     for (dup in 1:length(duplicates[[u]])){
       duplicates[[u]][dup] <- paste(u, "-", dup, sep = "")
     }
   }
   x <- unsplit(duplicates, x)
 }
 x
}
```

```
load_n_rename <- function(ds){
  load(paste("../outDATA/featuretables/", ds, "_ft.RData", sep = ""))
  row.names(sample_ft$data) <- sample_ft$sample_data[row.names(sample_ft$data), 
                                                     "Simplified Sample Name"]
  row.names(sample_ft$sample_data) <- sample_ft$sample_data$
                                                     "Simplified Sample Name"
    # Alphabetical order sample meta data, and make row order match in counts
  sample_ft$sample_data <- sample_ft$sample_data[
                              order(row.names(sample_ft$sample_data)),]
  sample_ft$data <- sample_ft$data[order(row.names(sample_ft$data)),]
  return(sample_ft)
}
```

```
ft_rel_abund_plot <- function(FeatureTable_object, Summarized_as, Dataset_name){
  fancy_abundance_plot <- FeatureTable_object %>% 
    map_samples(relative_abundance) %>% 
    plot(ylab = "Relative abundance", 
       legend_title = Summarized_as, 
       num_features = 10) +
    ggtitle(paste(Dataset_name, Summarized_as)) +
    scale_fill_viridis_d(option = "plasma") +
    theme(axis.text.x = element_text(angle = 80, hjust = 1, vjust = 1, size = 6),
          legend.text = element_text(size = 8))
  return(fancy_abundance_plot)
}
```

```
ft_replace_Z_clr <- function(x){
  clr_ft <- x$
    replace_zeros(use_cmultRepl = TRUE, 
                  method = "SQ", 
                  z.delete = FALSE,
                  z.warning = 1)$ 
    # z.delete previous default was FALSE, and setting z.delete = FALSE is
    # the only way to reproduce the plots from my methods. Allowing the 
    # current default will remove data and produce entirely different figures!
                                  clr()
  return(clr_ft)
}
```

```
# This function allows us to assign the shape to a parent feature of the feature
# for color. Ex. shape = fruit, bread, colors = apples, oranges, Rye, Ciabatta.
# I am not really a fan of the output when viewed in PCA plot by colorblindness
# simulator. Nice concept, but would need more separation of colors.
color_2_shape <- function(data, color_elem, shape_elem){
  sample_data <- data %>%
    select(c(color_elem, shape_elem))
  first_occur_color <- sample_data %>%
    subset(subset = !duplicated(sample_data[color_elem]),
          select = c(shape_elem, color_elem))
 row.names(first_occur_color) <- first_occur_color[,color_elem]
 first_occur_color <- first_occur_color[order(row.names(first_occur_color)),]
 first_occur_color <- first_occur_color %>% select(shape_elem)
 duplicates <- split(first_occur_color, first_occur_color)
 for (uniques in 1:length(duplicates)){
   duplicates[[uniques]] <- c(19, 17, 15, 25, 13, 7, 8, 3)[uniques]
   }
 shape_list <- unsplit(duplicates, first_occur_color)
 return(shape_list)
}
```

```
ft_clr_pca <- function(FeatureTable_object, Dataset_name, Summarized_as){
  
  x_pca <- FeatureTable_object$pca_biplot(use_biplotr = TRUE,
                                include_sample_data = TRUE,
                                arrows = FALSE) 
  
  colors_n_fills <- viridis::plasma(n=(length(unique(
    x_pca$plot_elem$data_for_ggplot$`Sample Description`))+1))
  colors_n_fills <- colors_n_fills[-length(colors_n_fills)]
  # Remove the yellow since it doesn't plot nicely on PCA's white background

  fancy_pca <- 
    # We are opting not to use the biplot object and make our own for more 
    # control over the points color/fill/and shape
    ggplot(x_pca$plot_elem$data_for_ggplot,
           aes(x = PC1,
               y = PC2)) +
    geom_point(aes(color = `Sample Description`,
                    fill = `Sample Description`, # Fill so shape 25 can be used
                    shape = `Sample Description`),
                size = 3) +
    # scale_color_viridis_d("Sample Description",
    #                       option = "plasma") +
    # scale_fill_viridis_d("Sample Description",
    #                       option = "plasma") +
    scale_color_manual(name = "Sample Description",
                       values = colors_n_fills) +
    scale_fill_manual(name = "Sample Description",
                       values = colors_n_fills) +
    scale_shape_manual(name = "Sample Description",
                      values = c(19, 17, 15, 25, 13, 7, 8, 3)) + #allows upto 8
                       # values = color_2_shape(
                       #   data = x_pca$plot_elem$data_for_ggplot,
                       #   color_elem = "Sample Description",
                       #   shape_elem = "Sample_type")) +
    coord_fixed(xlim = x_pca$plot_elem$xlimits,
                ylim = x_pca$plot_elem$ylimits) +
    ggtitle(paste(Dataset_name, Summarized_as)) +
    xlab(x_pca$plot_elem$xlabel) +
    ylab(x_pca$plot_elem$ylabel) +
    theme(panel.grid.major = element_blank(),
        panel.grid.minor = element_blank(),
        panel.background = element_blank(),
        axis.line = element_line(colour = "black"))

  return(fancy_pca)
}
```

##### Update Meta Data

```
datasets <- c("16S",
              "18S",
              "arrA",
              "dsrB",
              "mcrA",
              "narG",
              "nirK",
              "nrfA",
              "phoD",
              "phoX",
              "pmoA")
```

#### Relative Abundance

##### Collapse to Class

```
for (ds in datasets){
  # Previous featuretable object
  sample_ft <- load_n_rename(ds)

  # collapse to Class, remove controls, remove features only seen with controls
  rank_Class_ft <- sample_ft$
    collapse_features(by = "Class", keep_hierarchy = TRUE)$
    keep_samples(Sample_type != "Control")$
    keep_features(function(feature) sum(feature) > 0)

  # relative abundance plot with upto 15 Classes or 14 + Other
 assign(paste0("rel_abund_", ds),ft_rel_abund_plot(rank_Class_ft,
                                      Dataset_name = ds,
                                      Summarized_as = "Class"))

}
```

```
## Scale for fill is already present.
## Adding another scale for fill, which will replace the existing scale.
## Scale for fill is already present.
## Adding another scale for fill, which will replace the existing scale.
## Scale for fill is already present.
## Adding another scale for fill, which will replace the existing scale.
## Scale for fill is already present.
## Adding another scale for fill, which will replace the existing scale.
## Scale for fill is already present.
## Adding another scale for fill, which will replace the existing scale.
## num_features was one less than total number of features. There will be an 'Other' category, but it will only contain 1 feature!
## Scale for fill is already present.
## Adding another scale for fill, which will replace the existing scale.
## Scale for fill is already present.
## Adding another scale for fill, which will replace the existing scale.
## Scale for fill is already present.
## Adding another scale for fill, which will replace the existing scale.
## Scale for fill is already present.
## Adding another scale for fill, which will replace the existing scale.
## Scale for fill is already present.
## Adding another scale for fill, which will replace the existing scale.
## Scale for fill is already present.
## Adding another scale for fill, which will replace the existing scale.
```

```
plot_grid(rel_abund_18S, rel_abund_arrA, rel_abund_mcrA, rel_abund_narG, rel_abund_nrfA, rel_abund_phoD, rel_abund_phoX, rel_abund_pmoA, ncol = 2)
```

##### Collapse to 75% Similarity

```
for (ds in datasets){
  # Previous featuretable object
  sample_ft <- load_n_rename(ds)

  # collapse to cluster_75, remove controls, remove features only seen with controls
  rank_cluster_75_ft <- sample_ft$
    collapse_features(by = "cluster_75", keep_hierarchy = TRUE)$
    keep_samples(Sample_type != "Control")$
    keep_features(function(feature) sum(feature) > 0)

  # relative abundance plot with upto 15 cluster_75's or 14 + Other
 assign(paste0("rel_abund_", ds),ft_rel_abund_plot(rank_cluster_75_ft,
                                      Dataset_name = ds,
                                      Summarized_as = "75% Similarity"))

}
```

```
## Scale for fill is already present.
## Adding another scale for fill, which will replace the existing scale.
## Scale for fill is already present.
## Adding another scale for fill, which will replace the existing scale.
## Scale for fill is already present.
## Adding another scale for fill, which will replace the existing scale.
## Scale for fill is already present.
## Adding another scale for fill, which will replace the existing scale.
## Scale for fill is already present.
## Adding another scale for fill, which will replace the existing scale.
## Scale for fill is already present.
## Adding another scale for fill, which will replace the existing scale.
## Scale for fill is already present.
## Adding another scale for fill, which will replace the existing scale.
## Scale for fill is already present.
## Adding another scale for fill, which will replace the existing scale.
## Scale for fill is already present.
## Adding another scale for fill, which will replace the existing scale.
## Scale for fill is already present.
## Adding another scale for fill, which will replace the existing scale.
## Scale for fill is already present.
## Adding another scale for fill, which will replace the existing scale.
```

```
plot_grid(rel_abund_18S, rel_abund_arrA, rel_abund_mcrA, rel_abund_narG, rel_abund_nrfA, rel_abund_phoD, rel_abund_phoX, rel_abund_pmoA, ncol = 2)
```

#### PCA

##### ASV

```
for (ds in datasets){
  # Previous featuretable object
  sample_ft <- load_n_rename(ds)

  # remove controls, remove features only seen with controls
  ASV_ft <- sample_ft$
    keep_samples(Sample_type != "Control")$
    keep_features(function(feature) sum(feature) > 0)
  

  # zero replace and clr transform for PCA
  clr_ASV_ft <- ft_replace_Z_clr(ASV_ft)

  # PCA with coloring on "Sample Description"
 assign(paste0("pca_", ds),ft_clr_pca(clr_ASV_ft, 
                         Dataset_name = ds, 
                         Summarized_as = "ASV"))

}
```

```
## No. adjusted imputations:  179 
## No. adjusted imputations:  453 
## No. adjusted imputations:  525 
## No. adjusted imputations:  481 
## No. adjusted imputations:  625 
## No. adjusted imputations:  192 
## No. adjusted imputations:  377 
## No. adjusted imputations:  351 
## No. adjusted imputations:  569 
## No. adjusted imputations:  448 
## No. adjusted imputations:  1112
```

```
plot_grid(pca_18S, pca_arrA, pca_mcrA, pca_narG, pca_nrfA, pca_phoD, pca_phoX, pca_pmoA, ncol = 2)
```

##### Collapse to 95% Similarity

```
for (ds in datasets){
  # Previous featuretable object
  sample_ft <- load_n_rename(ds)

  # collapse to cluster_95, remove controls, remove features only seen with controls
  rank_cluster_95_ft <- sample_ft$
    collapse_features(by = "cluster_95", keep_hierarchy = TRUE)$
    keep_samples(Sample_type != "Control")$
    keep_features(function(feature) sum(feature) > 0)

  # zero replace and clr transform for PCA
  clr_cluster_95_ft <- ft_replace_Z_clr(rank_cluster_95_ft)

  # PCA with coloring on "Sample Description"
  assign(paste0("pca_", ds),ft_clr_pca(clr_cluster_95_ft,
                          Dataset_name = ds,
                          Summarized_as = "95% Similarity"))
}
```

```
## No. adjusted imputations:  140 
## No. adjusted imputations:  407 
## No. adjusted imputations:  653 
## No. adjusted imputations:  660 
## No. adjusted imputations:  734 
## No. adjusted imputations:  494 
## No. adjusted imputations:  676 
## No. adjusted imputations:  956 
## No. adjusted imputations:  597 
## No. adjusted imputations:  427 
## No. adjusted imputations:  740
```

```
plot_grid(pca_18S, pca_arrA, pca_mcrA, pca_narG, pca_nrfA, pca_phoD, pca_phoX, pca_pmoA, ncol = 2)
```

##### Collapse to Class

```
for (ds in datasets){
  # Previous featuretable object
  sample_ft <- load_n_rename(ds)

  # collapse to Class, remove controls, remove features only seen with controls
  rank_Class_ft <- sample_ft$
    collapse_features(by = "Class", keep_hierarchy = TRUE)$
    keep_samples(Sample_type != "Control")$
    keep_features(function(feature) sum(feature) > 0)
  
 # zero replace and clr transform for PCA
  clr_rank_Class_ft <- ft_replace_Z_clr(rank_Class_ft)

  # PCA with coloring on "Sample Description"
  assign(paste0("pca_", ds),ft_clr_pca(clr_rank_Class_ft,
                          Dataset_name = ds,
                          Summarized_as = "Class"))
}
```

```
## No. adjusted imputations:  9 
## No. adjusted imputations:  37 
## No. adjusted imputations:  7 
## No. adjusted imputations:  10 
## No. adjusted imputations:  17 
## No. adjusted imputations:  17 
## No. adjusted imputations:  16 
## No. adjusted imputations:  26 
## No. adjusted imputations:  23 
## No. adjusted imputations:  8 
## No. adjusted imputations:  19
```

```
plot_grid(pca_18S, pca_arrA, pca_mcrA, pca_narG, pca_nrfA, pca_phoD, pca_phoX, pca_pmoA, ncol = 2)
```

#### SessionInfo

```
sessionInfo()
```

```
## R version 4.4.2 (2024-10-31)
## Platform: x86_64-redhat-linux-gnu
## Running under: Fedora Linux 40 (Workstation Edition)
## 
## Matrix products: default
## BLAS/LAPACK: FlexiBLAS OPENBLAS-OPENMP;  LAPACK version 3.11.0
## 
## locale:
##  [1] LC_CTYPE=en_US.UTF-8       LC_NUMERIC=C              
##  [3] LC_TIME=en_US.UTF-8        LC_COLLATE=en_US.UTF-8    
##  [5] LC_MONETARY=en_US.UTF-8    LC_MESSAGES=en_US.UTF-8   
##  [7] LC_PAPER=en_US.UTF-8       LC_NAME=C                 
##  [9] LC_ADDRESS=C               LC_TELEPHONE=C            
## [11] LC_MEASUREMENT=en_US.UTF-8 LC_IDENTIFICATION=C       
## 
## time zone: America/New_York
## tzcode source: system (glibc)
## 
## attached base packages:
## [1] stats     graphics  grDevices utils     datasets  methods   base     
## 
## other attached packages:
##  [1] cowplot_1.1.3       viridis_0.6.5       viridisLite_0.4.2  
##  [4] featuretable_0.0.11 lubridate_1.9.3     forcats_1.0.0      
##  [7] stringr_1.5.1       dplyr_1.1.4         purrr_1.0.2        
## [10] readr_2.1.5         tidyr_1.3.1         tibble_3.2.1       
## [13] ggplot2_3.5.1       tidyverse_2.0.0     biplotr_0.0.13     
## [16] magrittr_2.0.3     
## 
## loaded via a namespace (and not attached):
##  [1] sass_0.4.9            utf8_1.2.4            generics_0.1.3       
##  [4] lattice_0.22-6        stringi_1.8.4         hms_1.1.3            
##  [7] digest_0.6.37         evaluate_1.0.1        grid_4.4.2           
## [10] timechange_0.3.0      fastmap_1.2.0         Matrix_1.7-1         
## [13] jsonlite_1.8.9        survival_3.7-0        zCompositions_1.5.0-4
## [16] gridExtra_2.3         fansi_1.0.6           scales_1.3.0         
## [19] truncnorm_1.0-9       jquerylib_0.1.4       cli_3.6.3            
## [22] rlang_1.1.4           splines_4.4.2         munsell_0.5.1        
## [25] withr_3.0.2           cachem_1.1.0          yaml_2.3.10          
## [28] tools_4.4.2           tzdb_0.4.0            colorspace_2.1-1     
## [31] NADA_1.6-1.1          vctrs_0.6.5           R6_2.5.1             
## [34] lifecycle_1.0.4       MASS_7.3-61           pkgconfig_2.0.3      
## [37] pillar_1.9.0          bslib_0.8.0           gtable_0.3.6         
## [40] glue_1.8.0            highr_0.11            xfun_0.49            
## [43] tidyselect_1.2.1      rstudioapi_0.17.1     knitr_1.48           
## [46] farver_2.1.2          htmltools_0.5.8.1     rmarkdown_2.29       
## [49] labeling_0.4.3        compiler_4.4.2
```
