## Supplementary material for "A Sample to Results Workflow for Compositional Analysis of Multiplexed Amplicon Sequencing Experiments": Markdown for diversity estimates

Appendix A Figures All Samples Alpha


### Appendix A Figures All Samples Alpha

###### Alexa Bennett

#### 2024-12-22

###### Load Packages

```
library(magrittr)
library(biplotr)
library(tidyverse)
```

```
## ── Attaching core tidyverse packages ──────────────────────── tidyverse 2.0.0 ──
## ✔ dplyr     1.1.4     ✔ readr     2.1.5
## ✔ forcats   1.0.0     ✔ stringr   1.5.1
## ✔ ggplot2   3.5.1     ✔ tibble    3.2.1
## ✔ lubridate 1.9.3     ✔ tidyr     1.3.1
## ✔ purrr     1.0.2     
## ── Conflicts ────────────────────────────────────────── tidyverse_conflicts() ──
## ✖ tidyr::extract()   masks magrittr::extract()
## ✖ dplyr::filter()    masks stats::filter()
## ✖ dplyr::lag()       masks stats::lag()
## ✖ purrr::set_names() masks magrittr::set_names()
## ℹ Use the conflicted package (<http://conflicted.r-lib.org/>) to force all conflicts to become errors
```

```
library(featuretable) # Keep after tidyverse
```

```
## 
## Attaching package: 'featuretable'
## 
## The following object is masked from 'package:dplyr':
## 
##     collapse
## 
## The following objects are masked from 'package:purrr':
## 
##     keep, map
## 
## The following object is masked from 'package:biplotr':
## 
##     pca_biplot
```

```
library(viridis)
```

```
## Loading required package: viridisLite
```

```
library(cowplot)
```

```
## 
## Attaching package: 'cowplot'
## 
## The following object is masked from 'package:lubridate':
## 
##     stamp
```

```
library(ggpubr)
```

```
## 
## Attaching package: 'ggpubr'
## 
## The following object is masked from 'package:cowplot':
## 
##     get_legend
```

##### Datasets

The rRNA and functional gene estimates viewed here are for Shannon
estimates with perturbation set to 0.0001.

```
datasets <- c("16S",
              "18S",
              "arrA",
              "dsrB",
              "mcrA",
              "narG",
              "nirK",
              "nrfA",
              "phoD",
              "phoX",
              "pmoA")
rRNA_datasets <- c("16S", "18S")
```

###### Make some functions to clean-up chunks

##### Sample Naming for Readability

```
ds <- "16S"
  load(paste("../outDATA/featuretables/", ds, "_ft.RData", sep = ""))

    simplified_naming <- sample_ft$sample_data %>% 
      dplyr::select(c("Simplified Sample Name", "Sample_type"))
      # sample names are currently row.names()
    print(simplified_naming)
```

```
##                            Simplified Sample Name Sample_type
## DESS-process-2020               Process Control-1     Control
## Freshwater-000390                FW-Silver Lake-1  Freshwater
## Freshwater-000391-tech-1         FW-Silver Lake-2  Freshwater
## Freshwater-000391-tech-2         FW-Silver Lake-3  Freshwater
## Freshwater-000392                FW-Silver Lake-4  Freshwater
## Freshwater-000395                FW-Silver Lake-5  Freshwater
## Freshwater-000396-tech-1         FW-Silver Lake-6  Freshwater
## Freshwater-000396-tech-2         FW-Silver Lake-7  Freshwater
## Freshwater-000697        FW-Coursey Pond LT-end-1  Freshwater
## Freshwater-000700                FW-Silver Lake-8  Freshwater
## Freshwater-040627160                FW-DE River-1  Freshwater
## Freshwater-120918327                FW-DE River-2  Freshwater
## Freshwater-Coursey-1     FW-Coursey Pond LT-beg-1  Freshwater
## Freshwater-Coursey-2     FW-Coursey Pond LT-beg-2  Freshwater
## Freshwater-Coursey-3        FW-Coursey Pond MOD-1  Freshwater
## Freshwater-Coursey-4        FW-Coursey Pond MOD-2  Freshwater
## Freshwater-DE-River-1-MM            FW-DE River-3  Freshwater
## Positive-control-2020          Positive Control-1     Control
## Sediment-091007078            Sediment-DE River-1    Sediment
## Sediment-120918326            Sediment-DE River-2    Sediment
## Soil-000698                Soil-Coursey Landing-1        Soil
## Soil-12252019-1-tech-1        Soil-Agricultural-1        Soil
## Soil-12252019-1-tech-2        Soil-Agricultural-2        Soil
## Soil-12252019-2-tech-1        Soil-Agricultural-3        Soil
## Soil-12252019-2-tech-2        Soil-Agricultural-4        Soil
## Soil-12252019-3-tech-1        Soil-Agricultural-5        Soil
## Soil-12252019-3-tech-2        Soil-Agricultural-6        Soil
```

```
load("../outDATA/divnet/all_datasets_divnet_etimates.RData")

summary_sample_ds_estimates["Sample_type"] <- sapply(
      summary_sample_ds_estimates$sample_names,
      function(x) simplified_naming$"Sample_type"[
        row.names(simplified_naming) == x])
                                               
summary_sample_ds_estimates["sample_names"] <- sapply(
      summary_sample_ds_estimates$sample_names,
      function(x) simplified_naming$"Simplified Sample Name"[
        row.names(simplified_naming) == x])
```

#### Alpha Diversity Quick View

The Shannon and Simpson estimates include the confidence intervals at
95% as “upper” and “lower” in addition to the estimate. Spearman’s
Correlation Coefficient will use the estimates for comparison since it
can not handle the confidence intervals. I selected a non-parametric
test here since the data was not normally distributed.

```
divnet_shannon <- summary_sample_ds_estimates %>%
    dplyr::filter(ds %in% c("arrA",
                            "mcrA",
                            "narG",
                            "nrfA",
                            "phoD",
                            "phoX",
                            "pmoA") &
           estimate_method == "shannon" &
           perturbation == 0.0001, 
           similarity %in% c("95")) 


shannon_plot <- ggplot(divnet_shannon, 
                       aes(x =sample_names, 
                           y = estimate,
                           color = Sample_type)) +
  # ggplot2 plot with confidence intervals
  geom_point() +
  coord_fixed(ylim = c(0,7)) +
  geom_errorbar(aes(ymin = lower, ymax = upper)) +
  scale_color_manual(name = "Sample Type",
                     values = c("#004488",
                                "#DDAA33",
                                "#BB5566")) +
  labs(
       y = "Shannon Estimate",
       x = "Sample Name") +
  theme_bw() +
  theme(axis.text.x = element_text(angle = 80, 
                                   hjust = 1,
                                   size = 6),
        legend.text = element_text(size = 8)) +
  facet_wrap(~ ds, ncol = 2)

print(shannon_plot)
```

#### SessionInfo

```
sessionInfo()
```

```
## R version 4.4.2 (2024-10-31)
## Platform: x86_64-redhat-linux-gnu
## Running under: Fedora Linux 40 (Workstation Edition)
## 
## Matrix products: default
## BLAS/LAPACK: FlexiBLAS OPENBLAS-OPENMP;  LAPACK version 3.11.0
## 
## locale:
##  [1] LC_CTYPE=en_US.UTF-8       LC_NUMERIC=C              
##  [3] LC_TIME=en_US.UTF-8        LC_COLLATE=en_US.UTF-8    
##  [5] LC_MONETARY=en_US.UTF-8    LC_MESSAGES=en_US.UTF-8   
##  [7] LC_PAPER=en_US.UTF-8       LC_NAME=C                 
##  [9] LC_ADDRESS=C               LC_TELEPHONE=C            
## [11] LC_MEASUREMENT=en_US.UTF-8 LC_IDENTIFICATION=C       
## 
## time zone: America/New_York
## tzcode source: system (glibc)
## 
## attached base packages:
## [1] stats     graphics  grDevices utils     datasets  methods   base     
## 
## other attached packages:
##  [1] ggpubr_0.6.0        cowplot_1.1.3       viridis_0.6.5      
##  [4] viridisLite_0.4.2   featuretable_0.0.11 lubridate_1.9.3    
##  [7] forcats_1.0.0       stringr_1.5.1       dplyr_1.1.4        
## [10] purrr_1.0.2         readr_2.1.5         tidyr_1.3.1        
## [13] tibble_3.2.1        ggplot2_3.5.1       tidyverse_2.0.0    
## [16] biplotr_0.0.13      magrittr_2.0.3     
## 
## loaded via a namespace (and not attached):
##  [1] sass_0.4.9        utf8_1.2.4        generics_0.1.3    rstatix_0.7.2    
##  [5] stringi_1.8.4     hms_1.1.3         digest_0.6.37     evaluate_1.0.1   
##  [9] grid_4.4.2        timechange_0.3.0  fastmap_1.2.0     jsonlite_1.8.9   
## [13] backports_1.5.0   Formula_1.2-5     gridExtra_2.3     fansi_1.0.6      
## [17] scales_1.3.0      jquerylib_0.1.4   abind_1.4-8       cli_3.6.3        
## [21] rlang_1.1.4       munsell_0.5.1     withr_3.0.2       cachem_1.1.0     
## [25] yaml_2.3.10       tools_4.4.2       tzdb_0.4.0        ggsignif_0.6.4   
## [29] colorspace_2.1-1  broom_1.0.7       vctrs_0.6.5       R6_2.5.1         
## [33] lifecycle_1.0.4   car_3.1-3         pkgconfig_2.0.3   pillar_1.9.0     
## [37] bslib_0.8.0       gtable_0.3.6      glue_1.8.0        highr_0.11       
## [41] xfun_0.49         tidyselect_1.2.1  rstudioapi_0.17.1 knitr_1.48       
## [45] farver_2.1.2      htmltools_0.5.8.1 labeling_0.4.3    carData_3.0-5    
## [49] rmarkdown_2.29    compiler_4.4.2
```
